# Cerebral oxygenation during locomotion is modulated by respiration

**DOI:** 10.1101/639419

**Authors:** Qingguang Zhang, Morgane Roche, Kyle W. Gheres, Emmanuelle Chaigneau, William D. Haselden, Serge Charpak, Patrick J. Drew

## Abstract

In the brain, increased neural activity is correlated with an increase of cerebral blood flow and increased tissue oxygenation. However, how cerebral oxygen dynamics are controlled in the behaving animals remains unclear. Here, we investigated to what extent the cerebral oxygenation varies during natural behaviors that change the whole-body homeostasis, specifically exercise. We measured oxygen levels in the cortex of awake, head-fixed mice during locomotion using polarography, spectroscopy, and two-photon phosphorescence lifetime measurements of oxygen sensors. We found that locomotion significantly and globally increases cerebral oxygenation, specifically in areas involved in locomotion, as well as in the frontal cortex and the olfactory bulb. The oxygenation increase persisted when neural activity and functional hyperemia were blocked, occurred both in the tissue and in arteries feeding the brain, and was tightly correlated with respiration rate and the phase of respiration cycle. Thus, respiration provides a dynamic pathway for modulating cerebral oxygenation.

---

An adequate oxygen supply is critical for proper brain function^1^, and deficiencies in tissue oxygen is a noted comorbidity in human diseases^2^ and aging^3^. For these reasons, there has been a great deal of interest in studying dynamics of cerebral oxygenation^4–9^. However, there is a gap in our understanding of how behavior, such as natural exercises like locomotion, affects cerebral oxygenation. In natural environments, animals and humans have evolved to spend a substantial portion of their waking hours locomoting^10^. As exercise is known to have a positive effect on brain health^11, 12^, a better understanding of the basic brain physiology accompanying the behaviors can give insight into how exercise can improve brain function. During movement, neuromodulator release and neural activity in many brain regions is elevated^13–20^, and there is an increase in cardiac output and respiratory rate. How these changes in local and systemic factors interact to control cerebral oxygenation is a fundamental question in brain physiology but is not well understood. Most cerebral oxygenation studies are performed in anesthetized animals^8, 9, 21–23^ (but see^4^), or non-invasively in humans. Anesthesia causes large disruptions of brain metabolism and neural activity^24^, and non-invasive human studies are impeded by technical issues, making accurate determination of any aspect of brain tissue oxygenation problematic.

Here we investigated how and by what mechanisms voluntary exercise impacts brain tissue oxygenation. We used intrinsic optical signal imaging^13, 25^, electrophysiology, Clark-type polarography^5, 6, 23^, and two-photon phosphorescent dye measurement^4, 8, 9^ to elucidate how vasodilation, neural activity, and systemic factors combine to generate changes in brain oxygenation. All experiments were performed in awake mice that were head-fixed on a spherical treadmill^13, 14^ or rotating disk^4^ that allowed them to voluntarily locomote. We found that cerebral oxygenation rose during locomotion in cortical regions that did not experience vasodilation, as well as when vasodilation was blocked. Oxygen levels increased in the arteries that supply the cortex during exercise, consistent with an increase in systemic oxygenation. Finally, we found that oxygen fluctuations were correlated with spontaneous and locomotion-evoked changes in respiration rate, as well as the phase of the respiration cycle, also consistent with a dynamic regulation in systemic oxygenation.

## Locomotion drives vasodilation in somatosensory, but not frontal cortex

We first assessed the spatial extent of cortical hemodynamic responses and their relationship to voluntary locomotion using intrinsic optical signal (IOS) imaging^13, 25^ (Fig. 1a). Imaging was done through a thin-skull window over the right-hemisphere (Fig. 1b). When the brain is illuminated with 530 nm light, reflectance decreases report dilations of arteries, capillaries and veins, which correspond with increases in cerebral blood volume (CBV). This reflectance change observed with IOS closely tracks measurements of vessel diameter made with two-photon microscopy^26^. The consistency with microscopic measurements of vessel diameter, combined with its very high signal-to-noise ratio^25^, and spatial resolution (less than 200 µm^27^), makes IOS suitable for detecting hemodynamic responses to locomotion. While neurally-evoked dilations initiate in the deeper layers of the cortex, the dilations propagate up the vascular tree to the surface arteries^28^, where they can be easily detectable with IOS. During locomotion, we observed region-specific changes in reflectance. There was a pronounced decrease in the reflectance (corresponding to an increase in CBV) in forelimb/hindlimb representation of the somatosensory cortex (FL/HL), while in frontal cortex (FC) there was no change, or a slight increase in reflectance (n = 11 mice, Fig. 1b, d). To better localize the area of decreased CBV, we used a smaller region of interest (ROI, 2 to 4 mm rostral and 0.5 to 2.5 mm lateral from bregma, ∼ 4 mm^2^) more rostral in FC than in our previous study^13^. We also assayed cerebral blood flow (CBF) using laser Doppler flowmetry, which will evaluate flow changes in a ∼1 mm^2^ area. The locomotion-evoked CBF showed similar spatial pattern of responses (n = 5 mice, Fig. 1c, d) as CBV. We quantified how locomotion affected CBV and CBF in two complimentary ways. We calculated the locomotion-triggered average, generated by aligning the IOS or laser Doppler signals to the onset of locomotion (see Methods) using only locomotion events ≥ 5 seconds in duration (Fig. 1d). We also calculated the hemodynamic response function (HRF)^25, 29^, which is the linear kernel relating locomotion events to observed changes in CBV and CBF (Supplementary Fig. 1). Both measures showed a decrease in CBV and CBF in the FC during locomotion (Fig. 1d, Supplementary Fig. 1). This shows that locomotion and the accompanying cardiovascular changes do not drive global increases in CBF/CBV, rather CBF/CBV increases are under local control. This lack of non-specific flow increase in the cortex during locomotion is likely because of autoregulation of the feeding arteries at the level of the circle of Willis and increased blood flow to the muscles^30^.

**Fig. 1.**
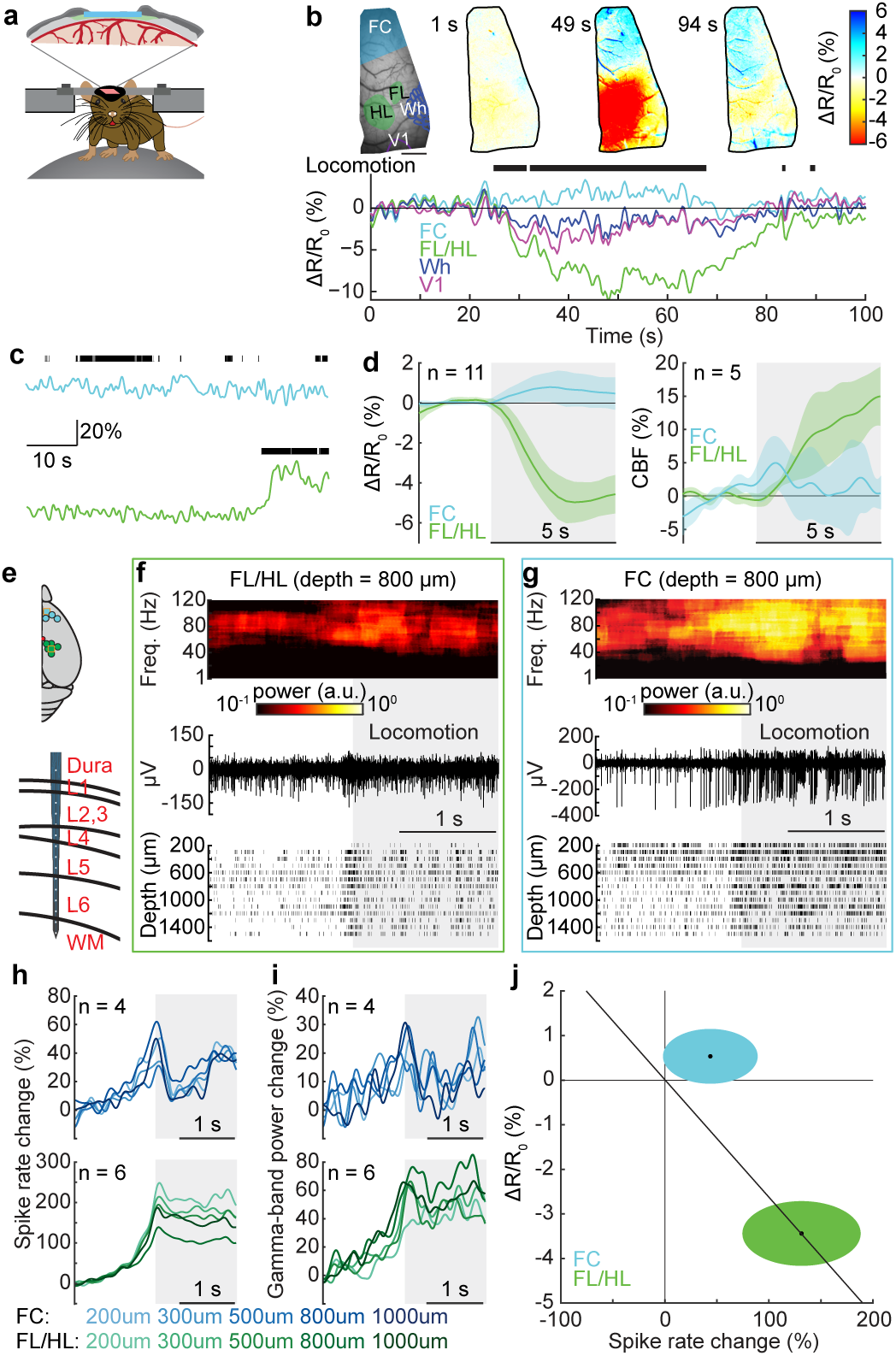
Locomotion drives cortical region-specific hemodynamic and neural responses across cortex. (**a**) Schematic of the experimental setup for IOS imaging. (**b**) Example data showing cerebral blood volume change during voluntary locomotion. Top left, an image of thin-skull window and corresponding anatomical reconstruction; scale bar = 1 mm. Top right, reflectance map before (1 s), during (49 s) and after (94 s) a voluntary locomotion event. Decreases in ΔR/R_0_ indicate increases in blood volume. Bottom, percentage change in reflectance (ΔR/R_0_) during locomotion events for each brain region. The black ticks denote locomotion events. FC, frontal cortex; FL/HL, forelimb/hindlimb representation of the somatosensory cortex; Wh, vibrissae cortex; V1, visual cortex. (**c**) Example trials showing locomotion-evoked changes of cerebral blood flow (CBF) in FC (top) and FL/HL (bottom) in the same animal. (**d**) Population average of locomotion-triggered average of CBV (n = 11 mice, left) and CBF (n = 5 mice, right) responses in both FL/HL (green) and FC (blue). (**e**) Top, schematic showing all laminar electrophysiology measurement sites in FC (n = 4 mice) and FL/HL (n = 6 mice). The squares indicate the measurement sites showing in (**f**) and (**g**). Bottom, schematic showing the layout of the electrodes and measurement depth. (**f**) Example trial showing the large increase in gamma-band power (top), raw signal (middle), and spike raster (bottom) during locomotion from a site 800 µm below the pia in FL/HL. Shaded area indicates the time of locomotion. (**g**) As in (**f**) but for FC. (**h**) Group average of locomotion evoked spike rate responses in both FC (top, n = 4 mice) and FL/HL (bottom, n = 6 mice). (**i**) As in (**h**) but for locomotion-evoked gamma-band LFP power responses. (**j**) Fractional change in the intrinsic signal, ΔR/R_0_, 2-5 s after the onset of locomotion plotted against spike rate change 0-2 s after the onset of locomotion in FL/HL (green ellipse) and FC (blue ellipse). For each ellipse, the radius along the vertical axis is the SD of ΔR/R_0_ across all animals (n = 11); the radius along the horizontal axis is the SD of spike rate across all animals (n = 4 for FC and n = 6 for FL/HL). The black dot in the center of each ellipse represents the average value of ΔR/R_0_ and spike rate response. The diagonal line shows the prediction of linear coupling.

To assess neural activity during locomotion, we measured local-field potential (LFP) and multi-unit activity (MUA) in a separate group of 7 mice (6 sites in FL/HL and 4 sites in FC) using multi-channel linear electrodes (Fig. 1e). We used electrophysiological measures of neural activity, as they are more sensitive than calcium indicators (which fail to detect about half of the spikes even under ideal conditions^31^), and do not disrupt normal neural activity as genetically encoded calcium indicators can do^32–34^. Since gamma-band (40-100 Hz) power in the LFP has been observed to be the strongest neural correlate of hemodynamic signals in rodents^25, 35^, primates^5, 36^ and humans^37^, and increases in gamma-band activity are also closely associated with the increases metabolic demand^38^, we quantified how locomotion affects neural activity by generating locomotion-triggered averages of gamma-band (40-100 Hz) power of LFP and spiking (see Methods). We observed that both the gamma-band power of LFP and spike rate increased during locomotion across all the layers in both FL/HL (Fig. 1f, h and i) and FC (Fig. 1g-i). The slow rise in neural activity a few hundred milliseconds before the onset of locomotion is due to low-pass filtering of the MUA signal (5 Hz, see Methods) and the windowing (1 second duration) required to estimate the LFP power^39^, as well as the ramping up of neural activity due to arousal changes seen before voluntary locomotion^40^. As optogenetic stimulation of fast spiking inhibitory neurons has been shown to induce large increases in blood oxygenation in the somatosensory cortex^41^, we sorted recorded spikes into fast spiking (FS, putatively inhibitory) and regular spiking (RS, putatively excitatory) spikes (see Methods). We found that FS and RS neurons exhibited a similar degree of rate increases during locomotion in both the FL/HL and FC areas (Supplementary Fig. 2).

Taken together, our results show that a short bout of locomotion increases neural activity, which is followed by an increase in CBV and CBF in FL/HL, and a small decrease in CBV and CBF in FC. Together with our previous work^13, 26^, these results suggest that the coupling between neural activity and hemodynamics are brain region-specific (Fig. 1j), as seen in many other neurovascular coupling studies in the cortex and other brain regions^42–45^. The lack of observed vasodilation in the FC is not due to a lack of sensitivity of our IOS imaging paradigm, as if the vasodilation in FC had the same relationship to neural activity as in the somatosensory cortex, we would expect to see a 2% decrease in the reflectance (Fig. 1j), which is easily detectable with our IOS setup^25^. As tissue oxygenation reflects the balance between oxygen supply and utilization^46^, we would expect that in FL/HL, the increased activity of the neurons will be more than matched by an increased blood supply, leading to an increase in tissue oxygenation. However, the increased neural activity in FC during locomotion will not be matched by an increase in the blood supply and should lead to a decrease in oxygenation in FC.

## Locomotion drives cortex-wide increases in brain tissue oxygenation

To test if the brain region-dependent differences in neurovascular coupling drove regional differences in brain oxygen dynamics during locomotion, we measured partial pressure of tissue oxygen (PtO_2_) in awake, behaving mice (n = 37 mice, 23 in FL/HL, and 14 in FC; 148.2 ± 28.3 minutes of recording per mouse) using Clark-type polarographic electrodes^47^ (Fig. 2a). Signals from these electrodes are similar to those obtained with BOLD fMRI^5, 6^, but with sub-second response time (Supplementary Fig. 3a), long-term stability (Supplementary Fig. 3b) and higher spatial resolution. We measured oxygen dynamics at different cortical depths by sequentially advancing the probe from the cortical surface into deeper layers. We observed a laminar-dependence of resting PtO_2_ in awake mice, with smaller oxygenation in surface layers and greater oxygenation in deeper layers in both FL/HL and FC (Supplementary Fig. 4a, b). Resting PtO_2_ was similar at each cortical depth in both FL/HL and FC (Supplementary Fig. 4a, b). These results, together with the observation that resting PtO_2_ is similar in somatosensory cortex and the olfactory bulb glomerular layer^4^, indicate that the spatial distribution of oxygen in the brain under normal (non-anesthetized) physiological condition is homogenous. Locomotion produces large, sustained dilation of arteries^48^ and increases in CBF and CBV^13, 26^ in the somatosensory cortex. These locomotion-induced dilations were not due to systemic effects, as they have been shown to be unaffected by drugs that do not cross the blood brain barrier that increase or decrease the heart rate^49^ and are blocked by the suppression of local neural activity^50^. The locomotion-induced dilations are comparable in magnitude to those elicited by episodic whisker stimulation^25^ which is known to elevate oxygenation, so one would expect increases in tissue oxygenation in FL/HL during locomotion. As anticipated, we observed increases in PtO_2_ during locomotion in FL/HL in all layers (Fig. 2b-d). Because the supply of blood to FC does not increase, but the neural activity does, one would expect a decline in tissue oxygenation during locomotion. Surprisingly, we also observed a very similar PtO_2_ increase in FC (Fig. 2b-d) to that observed in the FL/HL, despite small decreases in CBV or CBF, and an increase in neural activity. The elevation of PtO_2_ in FC during locomotion suggests that other factors can increase oxygenation in the brain.

**Fig. 2.**
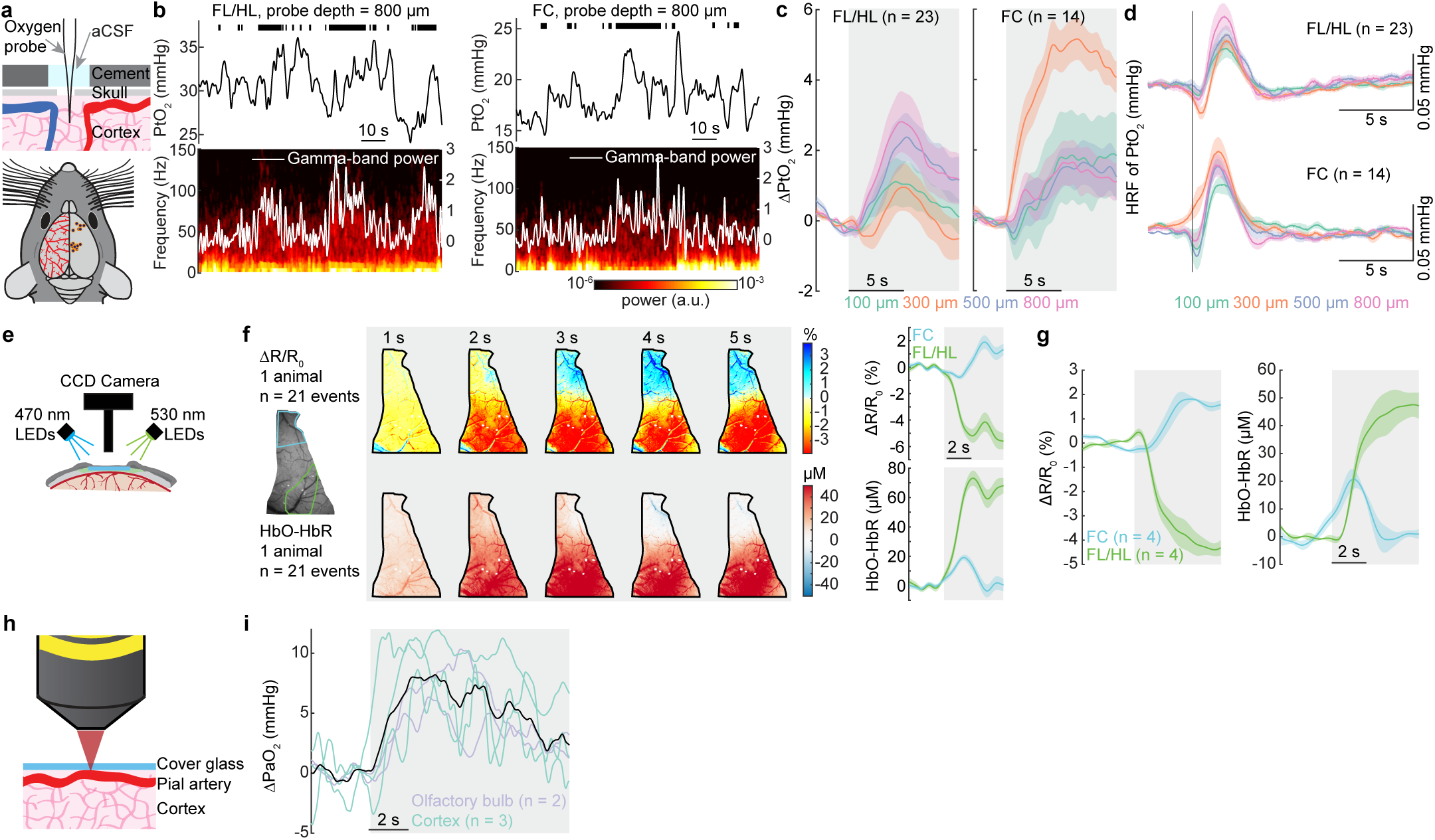
Cortex-wide increases in oxygenation during locomotion. (**a**) Top, a schematic showing the experimental setup. Bottom, measurement sites. (**b**) Example traces showing cortical tissue oxygenation (PtO_2_) responses to locomotion at sites 800 µm below brain surface in FL/HL (left) and FC (right). Top, black ticks denote binarized locomotion events; Middle, PtO_2_ responses to locomotion; Bottom, example of data showing spectrogram of LFP (white trace showing the gamma-band power). (**c**) Locomotion-evoked cortical tissue oxygenation increases (ΔPtO_2_) at all measured depths in both FL/HL (left, n = 23 mice) and FC (right, n = 14 mice). Gray shaded area indicates locomotion. Solid lines and shaded area denote mean ± standard error of the mean (SEM), respectively. (**d**) Hemodynamic response function (HRF) of tissue oxygenation at different depths in both FL/HL (top, n = 23 mice) and FC (bottom, n = 14 mice). Vertical black line showing the start of a brief impulse of locomotion. Data are shown as mean ± SEM. (**e**) Schematic showing the optical spectroscopy setup. (**f**) Left, example data showing spatial distribution of locomotion-evoked response of ΔR/R_0_ and difference between HbO and HbR (HbO-HbR) in an example mouse. Right, locomotion triggered average of ΔR/R_0_ and HbO-HbR for the same mouse in FC (blue) and FL/HL (green). (**g**) Group average of locomotion evoked response of ΔR/R_0_ and HbO-HbR in FC (n = 4 mice) and FL/HL (n = 4 mice). (**h**) Schematic showing the measurement of oxygen partial pressure in a cortical artery (PaO_2_) using two-photon phosphorescence lifetime microscopy (2PLM). (**i**) Locomotion induced PaO_2_ increases in 5 arteries (3 in the cortex (green) and 2 in the olfactory bulb (purple)) from a total of 4 mice. Mean response of all arteries is shown as a black line.

Polarographic probes provide measures of oxygen tension over a small region of brain tissue, and the response may be affected by the vasculature type and density^51–53^ surrounding the probe. To distinguish compartment-specific oxygen tension in the tissue, arterial and venous blood spaces, we then mapped the spatial distribution of locomotion-evoked brain oxygenation response using optical imaging spectroscopy^54, 55^ (Fig. 2e). Taking advantage of differences in the optical absorption spectra of oxyhemoglobin (HbO) and deoxyhemoglobin (HbR)^54, 55^, we collected reflectance images during rapid alternating green (530 nm) and blue (470 nm) illumination. Note that the spectroscopic measurements report oxygen concentrations in the red blood cells, while polarography reports average oxygen concentration in the tissue near the electrode. The oxygen levels in the tissue will differ from that in the blood somewhat due to the constraints of oxygen diffusion from the blood into the tissue and ongoing metabolic processes in the neurons and glial cells. Using the cerebral oxygenation index (HbO-HbR)^56^, the spectroscopic measures of hemoglobin oxygenation were similar to measurements from the tissue using polarographic probes: both methods yielded an increase in oxygenation during locomotion in both FC and FL/HL (Fig. 2f, g). These oxygenation changes persisted even when the heart rate increase associated with locomotion was pharmacologically blocked or occluded (Supplementary Fig. 5b, d, e), indicating they were not driven by the increased cardiac output during locomotion.

Moreover, the locomotion-induced elevation in oxygenation were present in the parenchyma, arterial and venous blood (Fig. 2f). As oxygen levels in the brain strongly depends on the arterial oxygen content^9^, we made direct measurements of oxygen partial pressure in the center of pial arteries (PaO_2_) using two-photon phosphorescence lifetime microscopy (2PLM, Fig. 2h)^4, 8, 9^, with a new phosphorescent probe (Oxyphor 2P) which has a very high brightness, improving measurement speed and imaging depth^57^. We asked if the oxygen levels increased in the center of the large pial arteries that supply blood to the brain. As the blood in these arteries will have minimal time to exchange oxygen in their transit through the heart and carotid artery to the brain, the oxygen levels in these arteries will track systemic oxygenation levels. We measured PaO_2_ in cortical and olfactory bulb arteries and found that PaO_2_ increased during locomotion (Fig. 2i). Taken together, these measurements are consistent with an increase in systemic blood oxygenation that leads to a brain-wide increase of oxygenation in the tissue and vascular compartments during locomotion. The increase in oxygenation accompanying the decrease in CBV and CBF in FC suggests that neurovascular coupling is not the only process controlling brain oxygenation^58^ during locomotion.

## Cortical oxygenation increases during locomotion even when vasodilation is blocked

Our observation that locomotion induced localized blood flow/volume increases, but cortical-wide increases in brain oxygenation, led us to hypothesize that the activity-dependent vasodilation may not be necessary for an increase in oxygenation. To test this, we pharmacologically blocked the glutamatergic and spiking activity by infusing/superfusing a cocktail of 6-cyano-7-nitroquinoxaline-2,3-dione (CNQX, 0.6 mM), (2R)-amino-5-phosphonopentanoic acid (AP5, 2.5 mM) and muscimol (10 mM) to suppress local neural activity. We first infused a cocktail of CNQX/AP5/muscimol via a cannula into FL/HL^25^, while concurrently monitoring neural activity, CBV and blood oxygenation (n = 4 mice, Fig. 3a). The cocktail infusion suppressed resting gamma-band (40-100 Hz) LFP power by 80 ± 12% and spiking activity by 82 ± 3% relative to vehicle infusions (Fig. 3b). Similarly, the standard deviation (SD) in gamma-band LFP power fluctuations during resting periods, an indicator of spontaneous neural activity levels, was decreased by 75 ± 18% in the gamma-band power and by 85 ± 6% in the MUA amplitude. To quantify the blood volume responses, we selected a semicircular region of interest (ROI) centered on the cannula and with a radius specified by the distance between the electrode and cannula (Fig. 3a), to ensure the ROI only included suppressed cortex^25^. Accompanying this neural activity blockade, baseline reflectance from the ROI increased (indicating decreased CBV, data not shown), and the locomotion-evoked decrease in reflectance (vasodilation) was almost completely suppressed (Fig. 3b-d), consistent with our previous study^25^ showing that intracerebral infusion of a cocktail of CNQX/AP5/muscimol suppressed sensory-evoked CBV increase. However, the block of neural activity was less effective during locomotion (Fig. 3b), likely due to the large increases of neural and modulatory drive to the cortex that occur during locomotion^17, 18, 59^. Nevertheless, this incomplete block of neural activity and block of vasodilation during locomotion is conducive for testing our hypothesis, as a complete block of vasodilation and an incomplete block of neural activity increases should lead to a *decrease* in oxygenation. However, if there is no oxygenation decrease, or the oxygenation increases, this would indicate that the oxygenation of the inflowing blood is elevated during locomotion. When locomotion-induced vasodilation was blocked, the locomotion-evoked increase in differences of oxy- and deoxygenated hemoglobin concentration (HbO-HbR) persisted, though the increase was smaller (Fig. 3b-d, Supplementary Fig. 5b, c, e). This increase was surprising, as we were able to completely block the locomotion-induced vasodilation, and there was still a small locomotion-induced increase in neural activity, which should result in a net decrease in oxygenation.

**Fig. 3.**
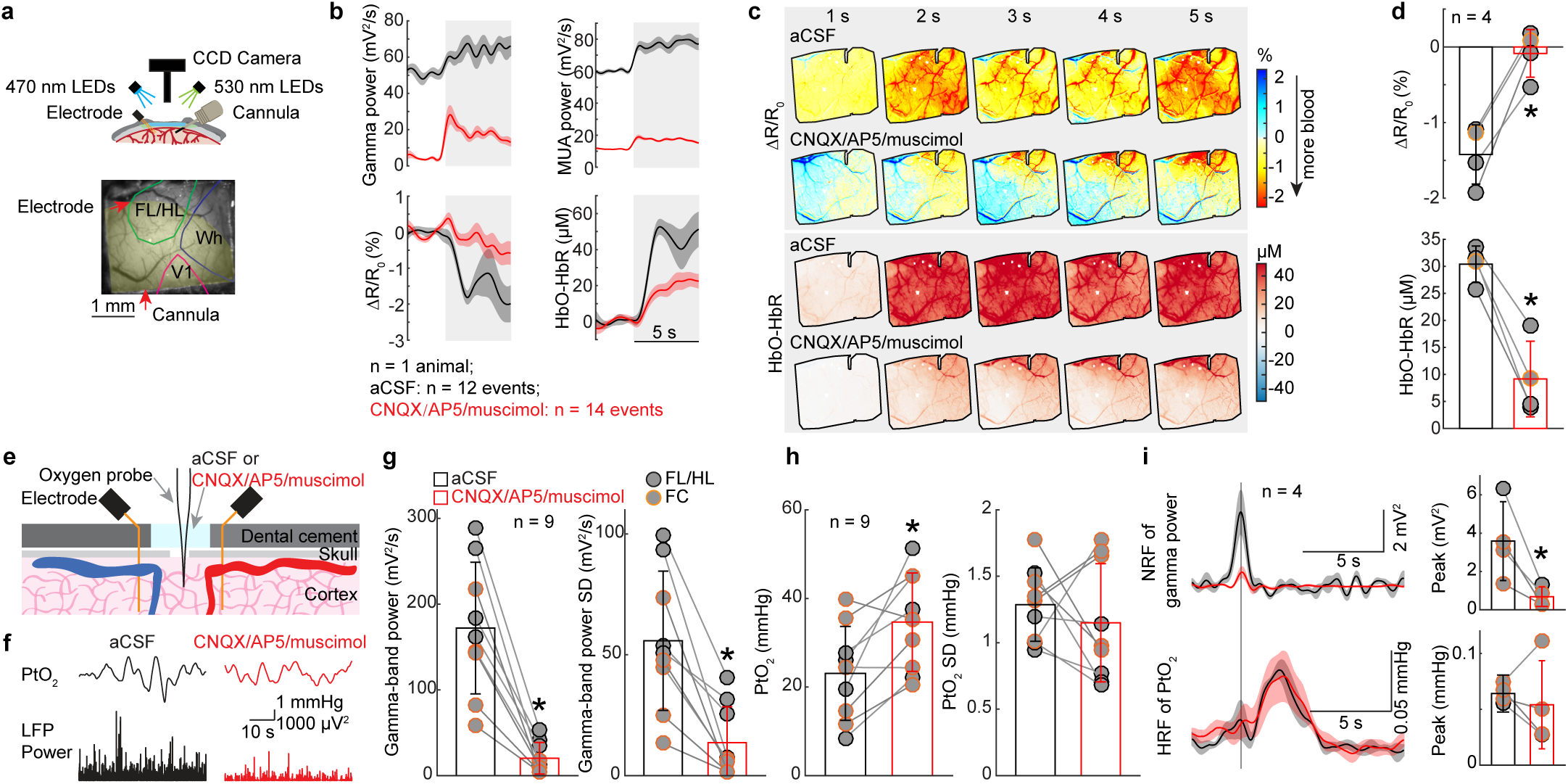
Locomotion-evoked cortical oxygenation increases persist when vasodilation is blocked. (**a**) Top, schematic of experimental setup for optical imaging spectroscopy measurement. Either aCSF or a cocktail of CNQX/AP5/muscimol was locally infused via a cannula. Bottom, an image of a polished thin-skull window with cannula and electrode implants. The yellow shaded area indicates the area affected by the drug infusion (i.e., region of interest for analysis). (**b**) Locomotion-evoked gamma-band (40-100 Hz) LFP power (top left), MUA power (top right), ΔR/R_0_ (bottom left) and difference between oxygenated and deoxygenated hemoglobin concentration (HbO-HbR, bottom right) in one representative mouse following aCSF (n = 12 locomotion events) and CNQX/AP5/muscimol (n = 14 locomotion events) infusion. Data was denoted as mean ± SEM. (**c**) Locomotion-evoked spatial distribution of ΔR/R_0_ (top) and HbO-HbR (bottom) for the same mouse shown in (**a**) and (**b**) following aCSF (n = 12 locomotion events) and CNQX/AP5/muscimol (n = 14 locomotion events) infusion. (**d**) Group average of locomotion-triggered ΔR/R_0_ (top, * paired *t*-test, t(3) = 7.4235, p = 0.0051) and HbO-HbR (bottom, * paired *t*-test, t(3) = 8.0007, p = 0.0041) signals after aCSF or CNQX/AP5/muscimol infusion in 4 mice. The orange circle denotes the mouse shown in (**b**) and (**c**). (**e**) Schematic of experimental setup for simultaneous tissue oxygenation and LFP measurements. CNQX (0.6 mM), AP5 (2.5 mM) and muscimol (10 mM) were added to aCSF bathing the craniotomy for 60-90 min, and recordings before and after the drug application were compared. (**f**) Example of resting PtO_2_ fluctuations (top) and resting gamma-band (40-100 Hz) LFP fluctuations (bottom) in the somatosensory cortex in a single mouse. (**g**) Comparison of spontaneous LFP activity (left, * Wilcoxon signed-rank test, p = 0.0039) and fluctuations (SD, right, * paired *t*-test, t(8) = 5.0246, p = 0.0010) before (black) and after (red) application of CNQX/AP5/muscimol in FL/HL (n = 4 mice, black circle) and FC (n = 5 mice, orange circle). (**h**) As (**g**) but for spontaneous PtO_2_ activity (left, * paired *t*-test, t(8) = 3.2712, p = 0.011) and fluctuations (SD, right, * paired *t*-test, t(8) = 0.7542, p = 0.4723). (**i**) Suppression of locomotion-evoked neural response does not affect locomotion-evoked ΔPtO_2_. Top left, neural response function (NRF) of gamma-band (40-100 Hz) power (n = 4 mice, 1 in FL/HL and 3 in FC) before (black) and after (red) application of CNQX/AP5/muscimol. Data are shown as mean ± SEM. Vertical black line indicates the start of a brief impulse of locomotion. Bottom left, as in top left but for HRF of PtO_2_. Top right, peak amplitude of NRF of gamma-band power before and after application of CNQX/AP5/muscimol (*paired *t*-test, one sided, t(3) = 3.4299, p = 0.0208). Bottom right, as in top right but for peak amplitude of HRF of PtO_2_ (paired *t*-test, t(3) = 0.5861, p = 0.599).

We further studied the effects of the suppressed vasodilation on oxygen responses in the tissue in a separate set of mice using polarographic electrodes (n = 9 mice, 5 in FC and 4 in FL/HL). We topically applied a cocktail of CNQX/AP5/muscimol to the cortex, while measuring spontaneous and locomotion-evoked neural activity and PtO_2_ in the superficial cortical layers (100-200 µm below the pia). The efficacy of the cocktail in suppressing neural activity was monitored with two electrodes spanning the oxygen measurement site^25, 35^ (Fig. 3e). Similar to intracortical infusions, superfusing the cocktail potently suppressed resting gamma-band LFP power by 89 ± 8% (Wilcoxon signed-rank test, p = 0.0039, Fig. 3g) and the SD by 77 ± 21% (paired *t*-test, t(8) = 5.02, p = 0.0010, Fig. 3g). Resting PtO_2_ increased by ∼70% following the suppression of neural activity and vasodilation (before: 23.09 ± 10.60 mmHg; after: 34.64 ± 11.11 mmHg; paired *t*-test, t(8) = 3.27, p = 0.011, Fig. 3h), consistent with neural signaling being a major component of metabolic demand^60^. To quantitatively assay locomotion-evoked oxygen and neural responses, we calculated the linear kernels (HRF) relating the PtO_2_ and the gamma-band power of LFP to locomotion. To ensure that vasodilation was blocked, we only analyzed those animals (n = 4 mice, 3 in FC and 1 in FL/HL) that showed >50% suppression of locomotion-evoked neural activity. In these animals, application of CNQX/AP5/muscimol reduced peak amplitude of gamma-band LFP neural response function (NRF) by 81 ± 8% (paired *t*-test, one sided, t(3) = 3.4299, p = 0.0208, Fig. 3i). If activity-dependent vasodilation is the only determinate of tissue oxygenation, we would expect the HRF of PtO_2_ shows profound reductions, since the vasodilation was blocked by the suppression of neural activity (Fig. 3b-d). However, the peak amplitude of PtO_2_ HRF was not changed (82 ± 51% of before cocktail application, paired *t*-test, t(3) = 0.5861, p = 0.599; Fig. 3i). Taken together, these results show that suppressing vasodilation does not block the locomotion-evoked oxygen increases.

## Respiration drives changes in cerebral and blood oxygenation

One possible driver of the increases in cerebral oxygenation is the increase in respiration during locomotion. Changes in respiration affect blood oxygen levels in the carotid artery^61, 62^ in anesthetized animals, and in humans, inhalation of 100% oxygen can elevate brain oxygen levels^63^. However, it is not known if normal fluctuations in respiration rate can impact cerebral oxygenation during normal behaviors. We tested whether respiration was correlated with oxygenation during locomotion by simultaneously measuring cortical tissue oxygenation and respiration (Fig. 4a). Locomotion was accompanied by a robust increase in respiratory rate (Fig. 4a, Supplementary Fig. 6), and fluctuations in respiratory rate on the time scale of seconds were linked to fluctuations in PtO_2_ (Fig. 4a). We quantified how well the fluctuations of respiratory rate and gamma-band (40-100 Hz) LFP power (which has been shown in previous studies to be the LFP band most correlated with vasodilation^25, 35^) correlated with the fluctuations in PtO_2_ by calculating the cross-correlation. During periods of rest, increases in gamma-band LFP power were correlated with decreased oxygenation (Fig. 4d, e), which was unexpected as gamma-band power increases during rest are correlated with vasodilation^25, 35, 64^. Because the decrease takes place with near zero time lag (Fig. 4d, e), it seems as though the dilation induced by spontaneous neural activity are insufficient relative to the metabolic demand. In contrast, respiration rate increases were correlated with increased oxygenation with a slight delay, consistent with the transit time of the blood from the lungs to the brain (Fig. 4b, c). When periods of locomotion were included, the correlation between gamma-band power and oxygenation was positive, suggesting that the coupling depends on animal’s state^64^ (Fig. 4d, e). The coupling between other frequency bands of the LFP and oxygen increases was negative (Supplementary Fig. 7), consistent with previous reports showing decreases in the power of these bands during voluntary locomotion^40^(also see Supplementary Fig. 8). Because cortical excitability and respiratory rate are correlated during locomotion (likely due to the reciprocal connections between respiratory and modulatory regions^65^), we sought to disentangle their respective contributions to cerebral oxygenation using partial coherence analysis^66^. We found that the coherence between respiratory rate and PtO_2_ was not due to the co-varying neural component (Supplementary Fig. 9b), nor was the coherence between gamma-band power and PtO_2_ affected by removing the respiratory rate contribution (Supplementary Fig. 9c). Thus, the partial coherence analysis indicates that respiration and neural activity (and likely vasodilation) affect tissue oxygenation independent of each other.

**Fig. 4.**
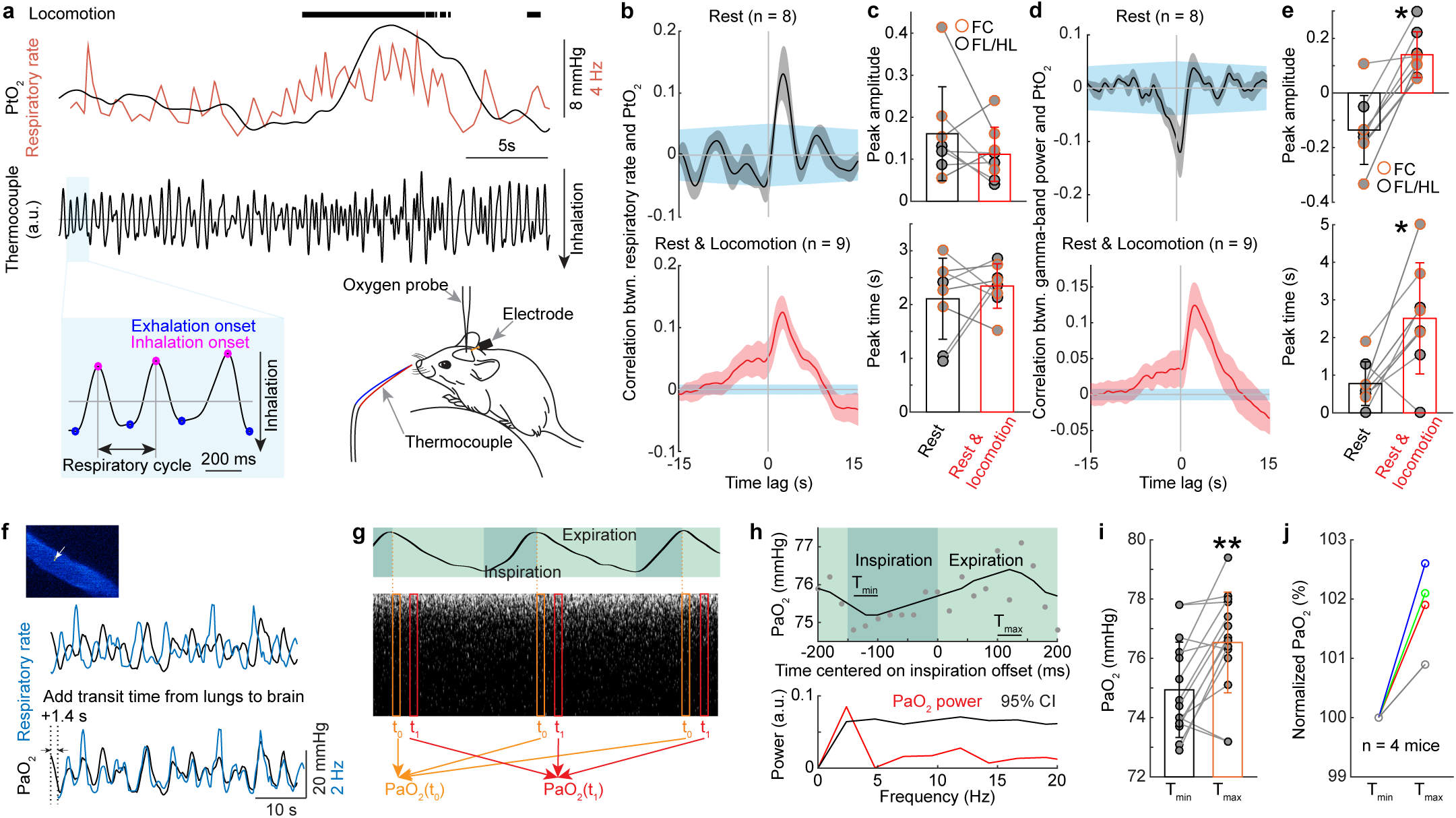
Respiration drives changes in cerebral tissue and arterial blood oxygenation. (**a**) Measuring respiration using a thermocouple. Top, example data showing tissue oxygenation (black trace) and raw respiratory rate (orange trace), during locomotion. Middle, thermocouple signal. Bottom left, expanded thermocouple signal showing of the detection of the onset of inspiratory (magenta dot) and expiratory phase (blue dot). Bottom right, schematic showing respiration measurement using a thermocouple. (**b**) Cross-correlation between PtO_2_ and respiratory rate signal from the thermocouple during periods of rest (top) and periods including rest and locomotion (bottom). The gray shaded region shows the population standard error of the mean. One mouse was excluded from resting correlation analysis as there were no resting segments long enough to meet the selection criteria. Blue shaded region shows 95% confidence intervals in cross-correlation obtained by shuffling the data. (**c**) Peak amplitude (top, Wilcoxon signed-rank test, p = 0.3125) and peak time delay (bottom, Wilcoxon signed-rank test, p = 0.7422) of cross-correlation between PtO_2_ and respiratory rate during periods of rest (black) and periods including rest and locomotion (red). (**d**) As (**b**) but for correlation between PtO_2_ and gamma-band power. (**e**) As (**c**) but for peak time (* paired *t*-test, t(7) = 6.1918, p < 0.001) and peak time delay (* Wilcoxon signed-rank test, p = 0.0234) of cross-correlation between PtO_2_ and gamma-band power. (**f**) Example data showing the temporal relation between respiratory rate (black) and oxygen tension (PaO_2_, blue) in the center of one artery (white arrow) in somatosensory cortex during periods of rest. The delay was due to transit time from lungs to brain. (**g**) Schematic showing the measurement of PaO_2_ fluctuations driven by the respiration cycle. Top, respiration signal and the segments of inspiration and expiration. Bottom, phosphorescent decay events are aligned to their position in the respiration cycle before being averaged into 20 ms bins. (**h**) PaO_2_ fluctuates within the respiratory cycle. Top, PaO_2_ change in one artery during the respiratory cycle at rest. PaO_2_ data (15 recordings with each of 50 seconds in duration) were aligned to the offset of inspiration. Each circle denotes averaged PaO_2_ over a short window (20 ms) aligned to a specific phase of respiration cycle and averaged over the 15 recordings. The solid curve shows the filtered data (first order binomial filter, 5 repetitions). T_min_ denotes the time period (40 ms) PaO_2_ reaches minimum. T_max_ denotes the time period (40 ms) PaO_2_ reaches maximum. Bottom, power spectrum of PaO_2_ (red) and 95% confidence interval (CI, black) given by randomizing the phase of the PaO_2_ signal. The PaO_2_ power at the respiratory frequency (∼2.5 Hz) is significantly greater than the 95% CI level. (**i**) PaO_2_ at maxima (T_max_) and minima (T_min_) for the 15 recordings from the artery shown in (**h**). ** p < 0.01, Wilcoxon signed-rank test. (**j**) Normalized PaO_2_ at minima (T_min_) and maxima (T_max_) for 4 vessels (3 in the cortex, and 1 in the olfactory bulb, n = 4 mice) with statistically significant PaO_2_ power spectrum peaks at the respiratory frequency. A total of 7 vessels were measured, and 4 out of 7 arteries showed significant peaks in the PaO_2_ power spectrum at the respiratory frequency.

The correlated fluctuations in respiratory rate and PtO_2_ suggests that the oxygen tension of arterial blood should also track the respiratory rate. To test this, we simultaneously monitored respiration and PaO_2_ in the pial arteries using 2PLM. In mice with irregular respiration, where respiratory rate transients of a few seconds occurred without locomotion, PaO_2_ followed respiration rate fluctuations (Fig. 4f), showing that changes in respiration rate can alter the oxygenation of the arterial blood entering the cortex.

We then asked if PaO_2_ tracked the phase of respiration, that is, whether the concentration of oxygen in the blood entering the brain fluctuated in phase with the inspiration-expiration cycle. This requires measuring PaO_2_ at rates high enough (> 5 Hz) to capture fluctuation in PaO_2_ due to respiration (nominally 2.5 Hz). As measurement of PaO_2_ with the 2PLM method is based on the *lifetime* of the phosphoresce decay of the dye, accurate quantification of the oxygen concentration requires averaging of decays^57^, which amounted to ∼3000 decays at our laser power (corresponding to ∼0.75 s of data), too slow to capture inspiration-expiration linked changes in PaO2. Therefore, we took advantage of the respiration cyclicality to collect sufficient amount of data. When the respiratory rate was very regular, the phosphorescence measures can be aligned and binned according to their place in the phase of the respiratory cycle (Fig. 4g), analogous to how erythrocyte-related transients can be detected in the capillaries^4, 8^, or analyzing the signal in the frequency domain. In a few animals with long bouts of highly regular respiration rate (average frequency 2.5 Hz, SD ≤ 0.6 Hz, average frequency/SD > 4), fluctuations of PaO_2_ tracked the respiratory cycle [4 out of 7 arteries (3 in the cortex, and 1 in the olfactory bulb) in 4 mice] (Fig. 4h-j). These arteries showed oscillations in PaO_2_ at the frequency of respiration that were significantly larger than would be expected by chance (reshuffling test, see Methods). This shows that the arterial blood flowing to the brain is not saturated at rest. It also shows that the oxygen tension in the blood tracks sub-second respiration dynamics, so increase in respiration can drive rapid increases in systemic blood oxygenation that will impact the brain oxygenation.

## Computational modeling indicates respiration contributes to tissue oxygenation

Using computer simulations, we then asked what the relative contributions of increased arterial oxygenation and vasodilation were to changes in PtO_2_. Recent work has shown that substantial oxygenation exchange occurs not only at capillaries, but also around the penetrating arteries in the cortex^7, 67^. To better understand how increase in blood oxygenation impact tissue oxygenation around arterioles, where the simple geometry of the vasculature allows us to better capture the dynamics of oxygenation changes due to vasodilation and systemic oxygenation changes, we created a Krogh cylinder model of a penetrating artery in the cortex^68^ (Fig. 5a). For this model, we used experimentally-determined quantities for the values of arterial oxygenation, and vessel diameter dynamics. We used published values for the cerebral metabolic rate of oxygen (CMRO_2_, Supplementary Table 1) for these simulations (Fig. 5b). The free parameters were chosen such that the tissue oxygenation predicted by the model matched our oxygen measurements in FC and FL/HL (Fig. 5c). Consistent with our data (Fig. 3h), the model also showed an increase in tissue oxygenation when neural activity (and metabolism) were suppressed (Fig. 5c). Moreover, using this model, we were able to tease out the relative contributions of vasodilation and increased arterial oxygenation to tissue oxygenation changes in both FC and FL/HL. In FL/HL, the large increase in CMRO_2_ during locomotion were counteracted by increases in arterial oxygenation due to vasodilation and increase in arterial oxygenation. In FC, the small increase in CMRO_2_ and vasoconstriction was totally offset by the increase in arterial oxygenation (Fig. 5d). These simulations show that respiration plays an important role in modulating tissue oxygenation. The increase in arterial oxygenation will also increase the oxygen tension in the tissue around the capillary bed^69^, though the actual changes will depend on the details of the capillary geometry and the movement of individual red blood cells, which is hard to capture without detailed anatomical models, and will depends on the details of flow dynamics. These simulations show that increased arterial oxygenation that accompanies increases in respiration can lead to increases in tissue oxygenation, even in brain regions showing vasoconstriction. Taken together, the experimental data and the simulation support the notion that increases in respiratory rate play an important role in regulating cerebral oxygenation.

**Fig. 5.**
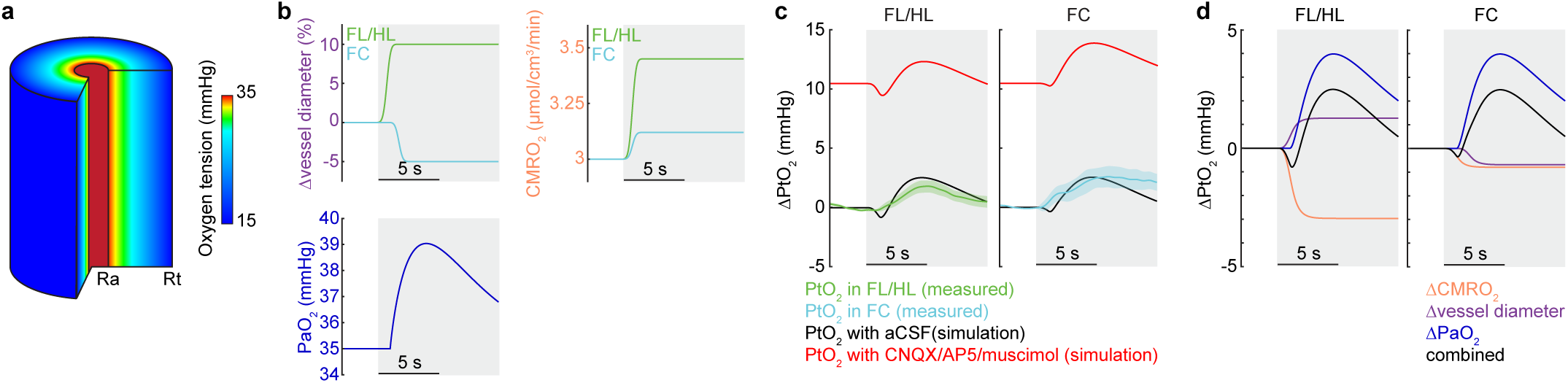
Tissue oxygenation during locomotion depends on the interplay of arterial oxygenation, CMRO_2_ and vasodilation. (**a**) Schematic showing the Krogh cylinder model of oxygen diffusion from a penetrating arteriole. An infinite tissue cylinder with radius R_t_ is supplied by an arteriole with radius R_a_. (**b**) Simulated locomotion induced changes in vessel diameters in FL/HL and FC (top left), cerebral metabolic rate of oxygen consumption (CMRO_2_) in FL/HL and FC (top right), and arteriole oxygen tension (PaO_2_, bottom). (**c**) Simulated effects of inhibiting neural activity using CNQX/AP5/muscimol on locomotion-evoked PtO_2_ change (ΔPtO_2_) in both FL/HL (left) and FC (right). Green and cyan shaded area denote one SEM of measured PtO_2_ change in FL/HL and FC, respectively. (**d**) Decomposition of locomotion-evoked oxygen changes in FL/HL (left) and FC (right). In both regions, changes in arterial oxygenation strongly influence tissue oxygenation.

## Discussion

We observed increases in cerebral tissue and blood oxygenation when respiration increased both at rest and during bouts of voluntary exercise. We also saw increases in tissue and blood oxygenation during locomotion when local neural activity was suppressed and vasodilation was blocked, conditions where we would expect a decrease in oxygenation. Note that while the changes in tissue and arterial oxygenation had similar dynamics (sustained increases in oxygen during locomotion), the oxygen increases measured spectroscopically were largest close to the onset of locomotion. This is likely because the spectroscopic imaging samples from arteries, capillaries, and veins. As the veins will be deoxygenated by the increased metabolic rate during periods of sustained neural activity, this will tend to reduce the measured oxygen change in the spectroscopic studies as compared to the polarography measurements, which primarily report tissue oxygen concentrations. Oxygen levels in the large arteries rose following increases in respiration both at rest and during exercise, and tracked the inspiration-expiration phase, showing that the oxygenation levels of the blood coming into the brain can be modulated by respiration both during rest and locomotion.

Respiration is not the only physiological change that accompanies exercise, and it bears considering other mechanisms that could account for the cerebral and arterial oxygenation changes seen here. Exercise causes large changes in cardiac output and blood pressure, and can be accompanied by changes in blood CO_2_ and lactate levels, but we think they are unlikely to be the cause of the nonspecific increase in cerebral oxygenation that we saw here. First, for the increases in cardiac output to raise global oxygenation in the cortex (independent of any changes in systemic oxygenation), it would need to drive an increase in cerebral blood flow. Our laser Doppler experiments show that blood flow does not rise in the frontal cortex, as they are likely buffered by autonomic regulation of the circle of Willis. Additionally, when heart rate and blood pressure increases during locomotion were blocked (with the beta blocker atenolol, which does not cross the blood brain barrier) or occluded (with the muscarinic receptor antagonist glycopyrrolate which also does not cross the blood brain barrier), there was no change in the locomotion-evoked CBV change (Supplementary Fig. 5, see also^49^). Therefore, systemic cardiac output increase cannot explain the increases in cerebral oxygenation seen during locomotion. Second, while CO_2_ is a strong vasodilator, and can drive increases in cerebral oxygenation under hypercapnia conditions by dilating blood vessels, rodents become hypocapnic during sustained exercise^70^. Exercise-evoked changes in CO_2_ would tend to cause cerebral vasoconstriction and would tend to drive a deoxygenation. Again, this mechanism could not drive the observed increase in blood and tissue oxygenation in the frontal cortex without corresponding flow increases and vasodilation. Sustained, high intensity exercise can cause increases in blood lactate over tens of minutes^71^, but there is no way that these lactate changes could drive changes in cerebral oxygenation seen in our experiments on the time scale of seconds. So, while many systemic variables change during voluntary locomotion, with the exception of increases in respiration rate, none would be able to increases the oxygen in the arteries or in the tissue within a few seconds of locomotion onset, nor could they explain the breathing cycle locked oscillations in the blood oxygenation or respiration-related fluctuations at rest. Thus, unless there is some heretofore unknown physiological process taking place during exercise, the most parsimonious explanation is that increases in respiration are the origin of the oxygenation increase in the brain observed here.

While our studies were performed in mice, there are respiration-driven fluctuations in the arterial blood of ungulates^61, 62^, suggesting it is a general property of mammals. While it is generally presumed that arterial blood is saturated in humans (but see^21^), arterial oxygen tension decreases substantially with age^72^ and acutely during sleep^73^. Respiration may play a more important role in cerebral oxygenation in humans than is currently appreciated, particularly as respiration rate is actively modulated during cognitive tasks^74, 75^. Respiration in humans is known to be increased following auditory or visual stimulation, and patterns of respiration differ from individual to individual, which might play a role in cerebral oxygen dynamics^76^.

The role of increased respiration in increasing brain oxygenation during behavior observed here is likely facilitated by the reciprocal connections between respiratory centers and the locus coeruleus^65, 77^ and other brain regions involved in arousal^78, 79^. Consistent with a tight interplay between respiration and metabolic demand in the brain, activation of the locus coeruleus, which will cause increases in alertness, and also causes concomitant increases in neural activity and blood flow in the cortex^80^. This tight interplay at the behavioral and anatomical levels between cortical arousal and respiration may help maintaining healthy oxygenation for optimal cortical function.

## Acknowledgements

This work was supported by a Scholar Award from the McKnight Endowment Fund for Neuroscience, and National Institutes of Health grants R01NS078168 and R01EB021703 to P.J.D., and by grants from the European Research Council (ERC-2013-AD6; 339513), the Agence Nationale de la Recherche (ANR/NSF 15-NEUC-0003-02), and the Fondation Leducq Transatlantic Networks of Excellence program (16CVD05) to S.C.. We thank A. K. Aydin for software to align locomotion, respiration and 2PLM oxygen tension data. Synthesis of the phosphorescent probe (Oxyphor 2P) was performed in the laboratory of Dr. Sergei Vinogradov (University of Pennsylvania) by Dr. Tatiana Esipova and supported by National Institutes of Health Grant R24NS092986 “Enabling widespread use of high resolution imaging of oxygen in the brain”.

## Author contributions

Q.Z. and P.J.D. designed the project. Q.Z. performed experiments and analyzed data using polarographic electrode, intrinsic optical imaging, optical imaging of spectroscopy, and laser Doppler. M.R. performed experiments and analyzed data using 2PLM. K.W.G. performed experiments and analyzed laminar electrophysiology data. E.C. analyzed 2PLM data. W.D.H. implemented the computational modeling. S.C. supervised experiments and data analysis using 2PLM. P.J.D. supervised experiments, modeling, data analysis, and preparation of the manuscript. Q.Z. and P.J.D. wrote the manuscript, with contributions from all authors.

## Materials and Methods

### Experimental design

Cerebral oxygenation, laminar electrophysiology, cerebral blood flow and volume data were acquired from separate groups of awake, behaving mice during voluntary locomotion. All experimental procedures were approved by the Pennsylvania State University and INSERM Institutional Animal Care and Use Committee guidelines.

### Animals

A total of 74 C57BL/6J mice (56 male and 18 female, 3-8 months old, 25-35 g, Jackson Laboratory) and 4 Thy1-GCaMP6f mice (3 male and 1 female, 3-12 months old, 25-35 g, Jackson Laboratory) were used. Recordings of laminar cortical tissue oxygenation were made from 37 mice [23 (13 male and 10 female) in the somatosensory cortex (FL/HL) and 14 (7 male and 7 female) in the frontal cortex (FC)] using Clark-type polarographic microelectrode. Simultaneous measurements of cortical tissue oxygenation using polarographic electrodes, respiration and local field potential were conducted in 9 mice [5 (4 male and 1 female) in FL/HL and 4 (2 male and 2 female) in FC]. Six of these mice were also used for laminar cortical tissue oxygenation measurements. Local field potential and spiking activity of different cortical layers were measured using laminar electrodes in a separate set of 7 male mice (4 in FC and 6 in FL/HL, 3 mice were measured in both FL/HL and FC simultaneously). Cerebral blood volume measurements using intrinsic optical signal imaging (with 530 nm illumination) were conducted in 11 male mice. Cerebral blood flow measurements using laser Doppler flowmetry were performed in 5 male mice. Tissue oxygenation measurements using spectroscopy (using alternating 470 nm and 530 nm illumination) were conducted in 11 male mice (4 mice were implanted with cannula and electrode). Oxygen measurements with 2PLM were conducted in adult Thy1-GCaMP6f (GP5.11) mice (n = 4). Mice were given food and water *ad libitum* and maintained on 12-hour (7:00–19:00) light/dark cycles. All experiments were conducted during the light period of the cycle.

### Surgery

With the exception of mice imaged with 2PLM, all surgeries were performed under isoflurane anesthesia (in oxygen, 5% for induction and 1.5-2% for maintenance). A custom-machined titanium head bolt was attached to the skull with cyanoacrylate glue (#32002, Vibra-tite). The head bolt was positioned along the midline and just posterior to the lambda cranial suture. Two self-tapping 3/32” #000 screws (J.I. Morris) were implanted into the skull contralateral to the measurement sites over the frontal lobe and parietal lobe. A stainless-steel wire (#792800, A-M Systems) was wrapped around the screw implanted in the frontal bone for use as an electrical ground for cortical tissue oxygenation and neural recordings. For cerebral blood flow (CBF) measurement using laser Doppler flowmetry (n = 5 mice), cerebral blood volume (CBV, n = 11 mice) measurement using intrinsic optical signal (IOS) imaging or brain oxygenation measurement using spectroscopy (n = 11 mice), a polished and reinforced thin-skull (PoRTS) window was made covering the right hemisphere as described previously^13, 25, 26, 49, 81^. For simultaneous measurement of tissue oxygenation and neural activity (n = 9 mice), we implanted two electrodes to measure LFP signals differentially. Electrodes were made from Teflon-coated tungsten wire (#795500, A-M Systems) with ∼1 mm insulation striped around the tip. The electrodes were inserted into the cortex to a depth of 800 µm at 45° angle along the rostral/caudal axis using a micromanipulator (MP-285, Sutter Instrument) through two small burr holes made in the skull. The two holes for the electrodes were made ∼1-1.5 mm apart to allow insertion of the oxygen probe between the two electrodes in following experiments. The holes were then sealed with cyanoacrylate glue. For spectroscopy imaging experiments with intracortical infusion (n = 4 mice), two small craniotomies were made at the edge of the thinned area of skull, and a cannula (dummy cannula: C315DCS; guide cannula: C315GS-4, Plastic One) was inserted into the upper layers of cortex at a 45° angle via one craniotomy. The stereotrode was placed 1.75 ± 0.5 mm away from the cannula through the other craniotomy. The screws, ground wire, electrodes and cannula were connected to the head-bolt via a midline suture using cyanoacrylate glue and black dental acrylic resin (#1530, Lang Dental Manufacturing Co.) to minimize skull movements. For tissue oxygenation (n = 37 mice) and laminar electrophysiology (n = 8 mice) experiments, the measurement sites were marked with ink and covered with a thin layer of cyanoacrylate glue. For oxygenation measurements using 2PLM, we used mice chronically implanted with a cranial window over FL/HL (n = 3 mice) or the olfactory bulb (n = 1 mice), using the protocol described previously^4^. Following the surgery, mice were then returned to their home cage for recovery for at least one week, and then started habituation on experimental apparatus.

### Habituation

Animals were gradually acclimated to head-fixation on a spherical treadmill^13, 24, 48^ or a rotating disk^4^ with one degree of freedom over at least three habituation sessions. The spherical treadmill was covered with nonabrasive anti-slip tape (McMaster-Carr) and attached to an optical rotary encoder (#E7PD-720-118, US Digital) to monitor locomotion. Mice were acclimated to head-fixation for ∼15 minutes during the first session and were head-fixed for longer durations (> 1 hour) in the subsequent sessions. Mice were monitored for any signs of stress during habituation. In all cases, the mice exhibited normal behaviors such as exploratory whisking and occasional grooming after being head-fixed. Heart-rate related fluctuations were detectable in the intrinsic optical signal^49^ and varied between 7 and 13 Hz for all mice after habituation, which is comparable to the mean heart rate (∼12 Hz) recorded telemetrically from mice in their home cage^82^. For oxygen measurements using 2PLM, a rotating disk treadmill was added to the cage a week prior to the surgery and restraint-habituation sessions started 3-4 days after surgery recovery. For habituation for 2PLM experiments, the animals were place head-fixed below the microscope and free to run on the treadmill. During each habituation session, a thermocouple (same as used for imaging) was placed close to the nostril in order to acclimate the mouse with its presence. Habituation sessions were performed 2-4 times per day over the course of one week, with the duration increasing from 5 min to 45 min.

### Physiological measurements

Data from all experiments were collected using custom software written in LabVIEW (version 2014, National Instruments).

#### Behavioral measurement

The treadmill movements were used to quantify the locomotion events of the mouse. The animal was also monitored using a webcam (Microsoft LifeCam Cinema®) as an additional behavioral measurement.

#### Cerebral tissue oxygenation measurement using polarographic electrode

On the day of measurement, the mouse was anesthetized with isoflurane (5% for induction and 2% for maintenance) for a short surgical procedure (∼20 min). A small (∼100 x 100 μm) craniotomy was made over the frontal cortex (1.0 to 3.0 mm rostral and 1.0 to 2.5 mm lateral from bregma) or the forelimb/hindlimb representation in the somatosensory cortex (0.5 to 1.0 mm caudal and 1.0 to 2.5 mm lateral from bregma), and dura was carefully removed (Fig. 2a). The craniotomy was then moistened with warm artificial cerebrospinal fluid (aCSF) and porcine gelatin (Vetspon). The mouse was then moved to and head-fixed on the spherical treadmill. Oxygen measurements started at least one hour after the mouse woke up from anesthesia to minimize the effects of anethesia^24, 83^.

Cerebral tissue oxygenation was recorded with a Clark-type oxygen microelectrode (OX-10, Unisense A/S, Aarhus, Denmark). A total of 9 probes were used in this study, with an average response time of 0.33 ± 0.11 seconds (n = 9 probes, Supplementary Fig. 3a and b). No compensation for the delay was performed. The oxygen electrodes were calibrated in air-saturated 0.9% sodium chloride (at 37°C) and oxygen-free standard solution [0.1M sodium hydroxide (SX0607H-6, Sigma-Aldrich) and 0.1M sodium ascorbate (A7631, Sigma-Aldrich) in 0.9% sodium chloride] before and after each experiment. The linear drift of the oxygen electrode signal (1.86±1.19% per hour, Supplementary Fig. 3c and d) was corrected by linearly interpolating between pre- and post-experiment calibrations. The oxygen electrode was connected to a high-impedance picoammeter (OXYMeter, Unisense A/S, Aarhus, Denmark), whose output signals were digitalized at 1000 Hz (PCI-6259, National Instruments). Current recordings were transformed to millimeters of mercury (mmHg) using the calibrations with air-saturated and oxygen-free solutions.

For oxygen polarography measurements, the oxygen microelectrode was positioned perpendicular to the brain surface and advanced into the cortex with a micromanipulator (MP-285, Sutter Instrument). The depth zero was defined as when the tip of the oxygen microelectrode touches the brain surface under visual inspection. The probe was then advanced to depth of 100, 300, 500 and 800 μm below the pia at the rate of 0.2 μm/step, and 30-40 min data were recorded for each depth. The tissue was allowed to recover for at least 5 min before the start of each recording.

In experiments investigating effects of suppressing vasodilation on cortical tissue oxygenation dynamics (Fig. 3e), a cocktail of ionotropic glutamate receptor antagonists 6-cyano-7-nitroquinoxaline-2,3-dione (CNQX, 0.6 mM), NMDA receptor antagonist (2R)-amino-5-phosphonopentanoic acid (AP5, 2.5 mM) and GABA_A_ receptor agonist muscimol (10 mM) were applied to suppress neural activity. All drugs were applied topically over the craniotomy and were allowed to diffuse into the cortical tissue for at least 90 min before the oxygen measurements. The efficacy of the CNQX/AP5/muscimol cocktail was monitored with simultaneously recorded neural activity. Neural data were amplified 1000x and filtered (0.1 – 10k Hz, DAM80, World Precision Instruments) and then sampled at 30k Hz (PCI-6259, National Instruments). The oxygen signal in these experiments was recorded at a depth of ∼100-200 μm.

At the end of the experiment, the mouse was deeply anesthetized, and a fiduciary mark was made by advancing an electrode (0.005” stainless steel wire, catalog #794800, A-M systems) into the brain with a micro-manipulator to mark the oxygen measurement site.

#### Respiration measurement using thermocouple

We conducted simultaneous respiration recordings in a subset of mice (n = 28) along with cortical oxygen measurements. Measurements of breathing were taken using 40-guage K-type thermocouples (TC-TT-K-40-36, Omega Engineering) placed near the mouse’s nose (∼ 1 mm), with care taken to not contact the whiskers. Data were amplified 2000x, filtered below 30 Hz (Model 440, Brownlee Precision), and sampled at 1000 Hz (PCI-6259, National Instruments). Downward and upward deflections in respiration recordings correspond to inspiratory and expiratory phases of the respiratory cycle, respectively (Fig. 4a). We identified the time of each expiratory peak in the entire record as the zero-crossing point of the first derivative of the thermocouple signal.

#### Laminar electrophysiology

Laminar electrophysiology recordings were performed in a separate set of mice (n = 7, Fig. 1e). On the day of measurement, the mouse was anesthetized using isoflurane (in oxygen, 5% for induction and 2% for maintenance). Two small (1×1 mm^2^) craniotomies were performed over the frontal cortex (1.0 to 2.5 mm rostral and 1.0 to 2.5 mm lateral from bregma) and FL/HL representation in the somatosensory cortex (0.5 to 1.0 mm caudal and 1.0 to 2.5 mm lateral from bregma) over the contralateral hemisphere (Fig. 1e), and the dura was carefully removed. The craniotomies were then moistened with warm saline and porcine gelatin (Vetspon). After this short surgical procedure (∼20 minutes), the mouse was then transferred to the treadmill where it was head-fixed. Measurements started at least one hour after the cessation of anesthesia^24, 83^.

Neural activity signals were recorded using two linear microelectrode arrays (A1×16-3mm-100-703-A16, NeuroNexus Technologies). The electrode array consisted of a single shank with 16 individual electrodes with 100 µm inter-electrode spacing. The signals were digitalized and streamed to SmartBox^TM^ via a SmartLink headstage (NeuroNexus Technologies). The arrays were positioned perpendicular to the cortical surface, one was in the forelimb/hindlimb representation in the somatosensory cortex and the other one was in the frontal cortex on the contralateral side. Recording depth was inferred from manipulator (MP-285, Sutter Instrument) recordings. The neural signals were filtered (0.1-10k Hz bandpass), sampled at 20k Hz using SmartBox 2.0 software (NeuroNexus Technologies).

#### Cerebral blood flow measurements using laser Doppler flowmetry

We measured cerebral blood flow responses to voluntary locomotion in a separate set of mice (n = 5) using laser Doppler flowmetry (OxyLab, Oxford Optronix)^49^. The probe was fixed 0.3 mm above the PoRTS window at a 45° angle. Data were sampled at 1000 Hz (PCI-6259, National Instruments).

#### Brain oxygen measurement using optical imaging of spectroscopy

We mapped the spatiotemporal dynamics of oxyhemoglobin and deoxyhemoglobin concentrations using their oxygen-dependent optical absorption spectra^54^. Reflectance images were collected during periods of green LED light illumination at 530 nm (equally absorbed by oxygenated and deoxygenated hemoglobin, M530L3, Thorlabs) or blue LED light illumination at 470 nm (absorbed more by oxygenated than deoxygenated hemoglobin, M470L3, Thorlabs). For these experiments, a CCD camera (Dalsa 1M60) was operated at 60 Hz with 4X4 binning (256 × 256 pixels), mounted with a VZM300i optical zoom lens (Edmund Optics). Green and blue reflectance data were converted to changes in oxy- and deoxyhemoglobin concentrations using the modified Beer-Lambert law with Monte Carlo-derived wavelength-dependent path length factors^54^. We used the cerebral oxygenation index^56^ (i.e., HbO-HbR) to quantify the change in oxygenation, as calculating the percentage change requires knowledge of the concentration of hemoglobin on a pixel-by-pixel basis, which is not feasible given the wide heterogeneity in the density of the cortical vasculature^51^.

In a subset of mice (n = 4), intracortical drug infusion were conducted via a cannula. Details of local infusion via a cannula were reported previously^25^. Briefly, mice were placed in the imaging setup, and we then acquired 40 min of imaging, neural and behavioral data with the dummy cannula in place. The dummy cannula was then slowly removed and replaced with an infusion cannula. The interface between the infusion cannula and the guide cannula was sealed with Kwik-Cast (World Precision Instruments). A cocktail of CNQX (0.6 mM)/AP5 (2.5 mM)/muscimol (10 mM), or L-NAME (100 µM), or aCSF was locally infused at a rate of 25 nL/min for a total volume of 500 nL. Drugs and vehicle controls were infused in a counterbalanced order. The efficacy of the CNQX/AP5/muscimol cocktail was monitored with simultaneously recorded neural activity using two tungsten electrodes. Neural data were amplified 1000x and digitally filtered (0.1-10k Hz, DAM80, World Precision Instruments) and then sampled at 30k Hz (PCI-6259, National Instruments). To verify that the dynamics observed after drug infusion were not due to changes of peripheral cardiovascular system^84, 85^, we also injected water, atenolol (2 mg/kg body weight) and glycopyrrolate (0.5 mg/kg body weight)^49^ intraparietal in the same mouse, and the hemodynamic response was measured described as above (Supplementary Fig. 5).

#### Brain oxygen measurement using two-photon phosphorescent lifetime microscopy

A complete description of 2PLM can be found in previous reports^4, 9, 86^. In brief, the oxygen sensor Oxyphor 2P^57^ was injected intravenously (final plasma concentration of 5 µM) under a brief isoflurane anesthesia (4%, < 3 min). The animals were allowed to recover for at least 1.5 h and then placed below the objective of a custom-built microscope. An acousto-optic modulator (AOM) was placed on the light path from Ti:Sapphire laser (Mira, Coherent; pulse width 250 fs, 76 MHz) to gate the 970 nm light excitation beam. Light was focused onto the center of pial arteries with a water-immersion objective (Olympus LUMFLN 60XW, NA 1.1) and collected emission was forwarded to a red-sensitive photomultiplier tube (PMT, R10699, Hamamatsu) after passing through a dichroic mirror (FF560-Di01, SEMROCK) and a band-pass filter (FF01-794/160, SEMROCK). PMT signals were amplified and sampled at 1.25 MHz by an acquisition card. PaO_2_ was estimated from the signal acquired during the AOM off-phase^9^, after discarding the first 5.6 µs (7 bins) following the end of the AOM on-phase. 200000 decays (50 s) were collected for each acquisition and 3000 decays were used for each lifetime measurement of PaO_2_. During the whole imaging session, respiration and locomotion were constantly monitored with the nasal thermocouple and a velocity encoder connected to the running wheel^87^.

#### Drugs

All drugs were purchased from Sigma-Aldrich except aCSF (#3525, Tocris) and sterile water (USP). Muscimol (M1523, 10 mM), CNQX (C127, 0.6 mM), AP5 (A5282, 2.5 mM) and L-NAME (N5751, 100 µM) were diluted in aCSF. Atenolol (A7655, 2 mg/kg body weight) and glycopyrrolate (SML0029, 0.5 mg/kg body weight) were diluted in sterile water. All drug solution was stored at −20°C and warmed up using a water bath (WB05A12E, PolyScience) immediately before application. Oxyphor 2P was kindly provided by Sergei Vinogradov.

### Histology

At the conclusion of the experiment, mice were deeply anesthetized with 5% isoflurane, transcardially perfused with heparinized saline, and then fixed with 4% paraformaldehyde. The brains were extracted and sunk in a 4% paraformaldehyde with 30% sucrose solution. The flattened cortices were sectioned tangentially (60 µm/section) using a freezing microtome and stained for the presence of cytochrome-oxidase^25, 52, 88, 89^. The anatomical locations of the oxygen measurement sites were then reconstructed using a combination of vascular images taken during surgery and the stained brain slices using Adobe Illustrator CS6 (Adobe Systems).

### Data analysis

All data analyses were performed in Matlab (R2015b, MathWorks) using custom code (by Q.Z., K.W.G. and P.J.D.).

#### Locomotion events identification

Locomotion events^13, 25, 48^ from the spherical treadmill were identified by first applying a low-pass filter (10 Hz, 5^th^ order Butterworth) to the velocity signal from the optical rotary encoder, and then comparing the absolute value of acceleration (first derivative of the velocity signal) to a threshold of 3 cm/s^2^. Periods of locomotion were categorized based on the binarized detection of the treadmill acceleration:

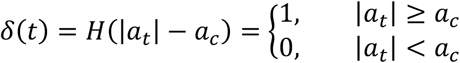

where *a_t_* is the acceleration at time t, and *a_c_* is the treadmill acceleration threshold.

#### Spontaneous and evoked activity

To characterize spontaneous (non-locomotion-evoked) activity, we defined “resting” periods as periods started 4 seconds after the end of previous locomotion event and lasting more than 10 seconds. Locomotion-evoked events were defined as segments with at least 3 seconds of resting prior to the onset of locomotion and followed by at least 5 seconds of locomotion. For oxygen measurements using polarographic electrode and two-photon phosphorescence lifetime microscopy, the locomotion segments need to be at least 10 seconds in duration.

#### Oxygen data preprocessing

Oxygen data from polarographic electrodes were first low-pass filtered (1 Hz, 5^th^ order Butterworth). The oxygen data were then down-sampled to 30 Hz to align with binarized locomotion events for calculation of locomotion-triggered average and hemodynamic response function.

#### Laminar neural activity

The neural signal was first digital filtered to obtain the local field potential (LFP, 0.1-300 Hz, 5^th^ order Butterworth) and multiunit activity (MUA, 300-3000 Hz, 5^th^ order Butterworth)^13, 25^. Time-frequency analysis of LFP signal was conducted using multi-taper techniques (Chronux toolbox version 2.11, http://chronux.org/)^90^. The power spectrum was estimated on a 1 second window with ∼1 Hz bandwidth averaged over nine tapers. MUA signals were low-pass filtered (5 Hz, Bessel filter). The locomotion-evoked LFP power spectrum was converted into relative power spectrum by normalizing to the 3 second resting period prior to the onset of locomotion. Spike rate was obtained by counting the numbers of events that exceed an amplitude threshold (three standard deviations above background) in each 1 millisecond bin.

#### Spike sorting

Sortable spike waveforms were extracted from MUA recordings using spike times identified from threshold crossings at four standard deviations of the mean. Spike waveforms were interpolated using a cubic spline function (MATLAB function: interp1) and were normalized by the amplitude of the peak of the action potential. We classified waveforms as fast spiking (FS) or regular spiking (RS) neurons based on the peak-to-trough-duration of the normalized waveform of each spike. Peak-to-trough times of all spikes across all layers were binned at 0.05 ms intervals (the minimal temporal resolution at 20kHz sampling rate). A histogram of peak-to-trough times was fitted as a sum of two Gaussian distributions (Supplementary Fig. 2a, f), and a receiver operator characteristic curve was used to segregate spikes in a given bin as either FS or RS waveforms using a 95% probability of belonging to a group as the inclusion threshold. Spikes not reaching the inclusion threshold for either group were not included in the analysis. Fast spiking (FS) waveforms (Supplementary Fig. 2b, g) were characterized by short durations between action potential peak and peak of hyperpolarization, peak-to-trough-duration as described previously^40, 91^. We characterized the RS and FS activity across different cortical layers during both resting and locomotion periods. To directly compare locomotion-related changes between FS and RS neurons, we calculated the percentage change of FS (ΔFS) and RS (ΔRS) spike rates (Supplementary Fig. 2c-e and h-j), which normalizes for absolute rate differences.

#### Calculation of hemodynamic response function and neural response function

We considered the neurovascular relationship to be a linear, time-invariant system^29, 92, 93^. To provide a model-free approach to assess the relationship between laminar tissue oxygenation and laminar neural activity, the hemodynamic response function (HRF) and neural response function (NRF) were calculated by deconvoluting tissue oxygenation signal, neural activity signal or respiratory rate signal to locomotion events, respectively, using the following equation:

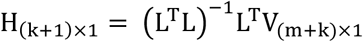

H is the HRF or NRF, V is the tissue oxygenation signal or neural activity signal. L is a Toeplitz matrix of size (m+k) x (k+1) containing measurements of locomotion events (n):

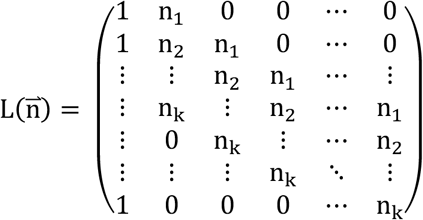

#### Cross-correlation analysis

Cross-correlation analysis was performed between simultaneously recorded neural/respiration and oxygen signals to quantify the relationship between fluctuations. For spontaneous correlations, only periods of rest lasting more than 30 seconds were used, with a four-seconds buffer at the end any locomotion event. We also calculated the correlations using all the data including periods with locomotion. To check the spatiotemporal distribution of the correlation, we calculated cross-correlogram between PtO_2_ and LFP power in each frequency band (Supplementary Fig. 7). Briefly, LFP signals were separated into frequency bands (∼1 Hz resolution with a range of 0.1-150 Hz) by calculating the spectrogram (mtspecgramc, Chronux toolbox)^90^, and then we calculate the temporal cross-correlation between power in each frequency band and the oxygen concentration (xcorr, MATLAB). Positive delays denote the neural signal lagging the oxygen signal. The oxygen tension and neural activity were both low-pass filtered below 1 Hz before calculating the cross-correlation. The temporal cross-correlation between respiratory rate and oxygen signals was also calculated over a similar interval (xcorr, MATLAB). Statistical significance of the correlation was computed using bootstrap resampling^94^ from 1000 reshuffled trials.

#### Arterial oxygen tension changes during the respiration cycle

To evaluate the arterial oxygen tension change within the respiration cycle, we selected oxygen measurements during periods with regular respiratory rate (average frequency 2.5 Hz, SD ≤ 0.6 Hz). The phosphorescent decays were aligned according to their place in the phase of the respiratory cycle (**Fig. 4** g). To further determine whether the fluctuations of oxygen tension was induced by respiration, we calculated the power spectrum of arterial oxygen tension, and determined the peak frequency in the power spectrum. Statistical significance of this peak was calculated by reshuffling the arterial oxygen measurements^94^, and the 95% confidence interval was calculated using 10000 reshuffled trials.

#### Ordinary coherence and partial coherence

We used coherence analysis^95^ to reveal correlated oscillations and deduce functional coupling among different signals. The ordinary coherence between two signals x and y are defined as

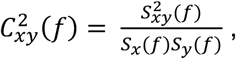

where *S*_*x*_(*f*) and *S*_*y*_(*f*) are the auto-spectra of the signals, and *S*_*xy*_(*f*) is the cross-spectrum. For ordinary coherence analysis between two signals (x and y), highly coherent oscillations can occur if they are functionally connected or because they share a common input. To differentiate between these possibilities, we also computed the partial coherence, i.e., coherence between two signals (x and y) after the removal of the components from each signal that are predictable from the third signal (z). The partial coherence function measuring the relationship of x and y at frequency f after removal of z is defined as

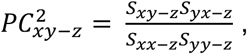

where *S*_*xx*-*z*_ and *S*_*yy*-*z*_ is the auto-spectra associated with the residual part of x and y after removing the part coherent with z, respectively. *S*_*xy*-*z*_ is the cross-spectrum between the residual part of x and y after removing the part coherent with z. If all the networks are connected, partial coherence will be between zero and the level of the ordinary coherence. If the connection behaves in an asymmetric manner, i.e., signal z affects x and y differentially, the coherence between two signals may increase after partialization (Supplementary Fig. 9a).

### Statistical analysis

Statistical analysis was performed using Matlab (R2015b, Mathworks). All summary data were reported as the mean ± standard deviation (SD) unless stated otherwise. Normality of the samples were tested before statistical testing using Anderson-Darling test (adtest). For comparison of multiple populations, the assumption of equal variance for parametric statistical method was also tested (vartest2 and vartestn). If criteria of normality and equal variance were not met, parametric tests (t test, one-way ANOVA) were replaced with a nonparametric method (Mann-Whiteney U-test, Wilcoxon signed-rank test, Kruskal-Wallis ANOVA). All *p* values were Bonferroni corrected for multiple comparisons. Significance was accepted at *p* < 0.05.

### Computational modelling

We simulated oxygen diffusion from a penetrating arteriole using the Krogh cylinder model^68^ using COMSOL (COMSOL Inc.). The concentration of oxygen([O_2_]) at any point in space was given by the equations:

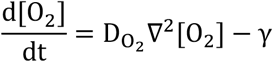

where D_o_2__ is the diffusion coefficient for oxygen in tissue (2800 µm^2^s^-1^) and γ is the cerebral metabolic rate of oxygen consumption (CMRO_2_) in the tissue. Resting arterial oxygen tension (PaO_2_) in the penetrating vessels was taken to be 35 mmHg^96^. The arterial and tissue radius was assumed to be 9 µm and 50 µm, respectively^68, 97^, with a periodic boundary condition beyond the tissue cylinder with a radius Rt. The oxygen consumption rate was uniform outside the arteriole, with a resting CMRO_2_ of 3 µmole/cm^3^/min^98^. The model was initialized at steady state. We assumed that locomotion induced a 15% increase in CMRO_2_ in FL/HL and a 4% increase in FC (proportionally scaled based on our neural recordings in Fig. 1f-i). We took the optical reflectance changes in FL/HL and FC to be 10% dilation and 5% constriction in vessel diameter, respectively, based on the measured relationship between reflectance and arteriole diameter in our previous study^26^. As ∼75% of neural tissue oxygen consumption is activity dependent^60, 99^, we simulated effects of CNQX/AP5/muscimol application by reducing the neuronally dependent portion of CMRO_2_ by 82%, yielding a CMRO_2_ of 1.2 µmole/cm^3^/min. Details of the model parameters are shown in Supplementary Table 1.

## Supplemental Information

**Supplementary Table 1.** Model parameters.

| Model parameter | Description | Value | Source |
| --- | --- | --- | --- |
| DO<sub>2</sub> | Oxygen diffusivity | 2800 μm<sup>2</sup>/s | 100 |
| R<sub>vessel</sub> | vessel radius | 9 μm |  |
| R<sub>t</sub> | Radius of tissue cylinder | 50 μm | 68,97 |
| CMRO<sub>2, baseline</sub> | Resting CMRO<sub>2</sub> | 3 μmole/cm<sup>3</sup>/min | 98 |
| CMRO<sub>2, MAC</sub> | CMRO<sub>2</sub> after CNQX/AP5/muscimol application | 1.2 μmole/cm<sup>3</sup>/min |  |
| PO<sub>2, baseline</sub> | Resting arterial PO<sub>2</sub> | 35 mmHg | 96 |
| ρ | solubility coefficient for O<sub>2</sub> | 1.39 μM/mmHg | 68,96,101 |
| Locomotion-evoked dynamics |  |  |  |
| $PO_{2, art} = \begin{cases} PO_{2, baseline} & \text{if } t < \tau \\ PO_{2, baseline} \left( A \frac{(t - \tau)^{\alpha - 1} \beta^\alpha e^{-\beta(t - \tau)}}{\Gamma(\alpha)} + 1 \right) & \text{if } t \geq \tau \end{cases}$ | | | |
| A |  | 1 |  |
| α |  | 1.9 |  |
| β |  | 0.3 |  |
| τ (time shift) |  | 1 s | 102 |
| Locomotion-evoked change of vessel radius |  |  |  |
| FL/HL |  | +10% (+0.9 μm) |  |
| FC |  | -5% (-0.45 μm) |  |
| CNQX/AP5/muscimol |  | -20% (-1.8 μm) |  |
| Locomotion-evoked change of CMRO<sub>2</sub> |  |  |  |
| FL/HL |  | +15% (+0.45 μmole/cm<sup>3</sup>/min) | 103-105 |
| FC |  | +4% (+0.12 μmole/cm<sup>3</sup>/min) |  |

**Supplementary Fig. 1.**
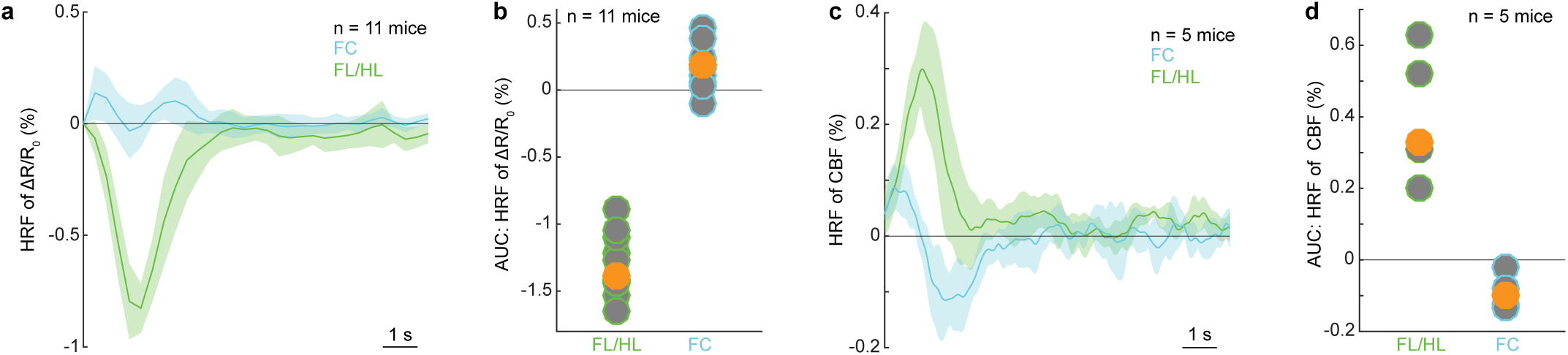
Cerebral blood flow (CBF) and volume (CBV) hemodynamic response functions. Related to **Fig. 1**a-d. (**a**) Hemodynamic response function (HRF) of reflectance change (ΔR/R_0_) in FL/HL (green) and FC (blue). (**b**) Integrated area under the curve (AUC) for HRF of (ΔR/R_0_) from 0 to 3 seconds. Each circle represents the HRF from a single mouse. The orange circle represents population median. AUC of the CBV HRF was less than zero in FL/HL (Wilcoxon signed-rank test, p < 0.0001) and greater than zero in FC (Wilcoxon signed-rank test, p = 0.002), indicating a dilation and constriction, respectively. (**c**) As in (**a**) but for cerebral blood flow (CBF). (**d**) As in (**b**) but for cerebral blood flow. AUC of HRF is greater than zero in FL/HL (Wilcoxon signed-rank test, p = 0.03) and less than zero in FC (Wilcoxon signed-rank test, p = 0.03), indicating increased and decreased flow, respectively.

**Supplementary Fig. 2.**
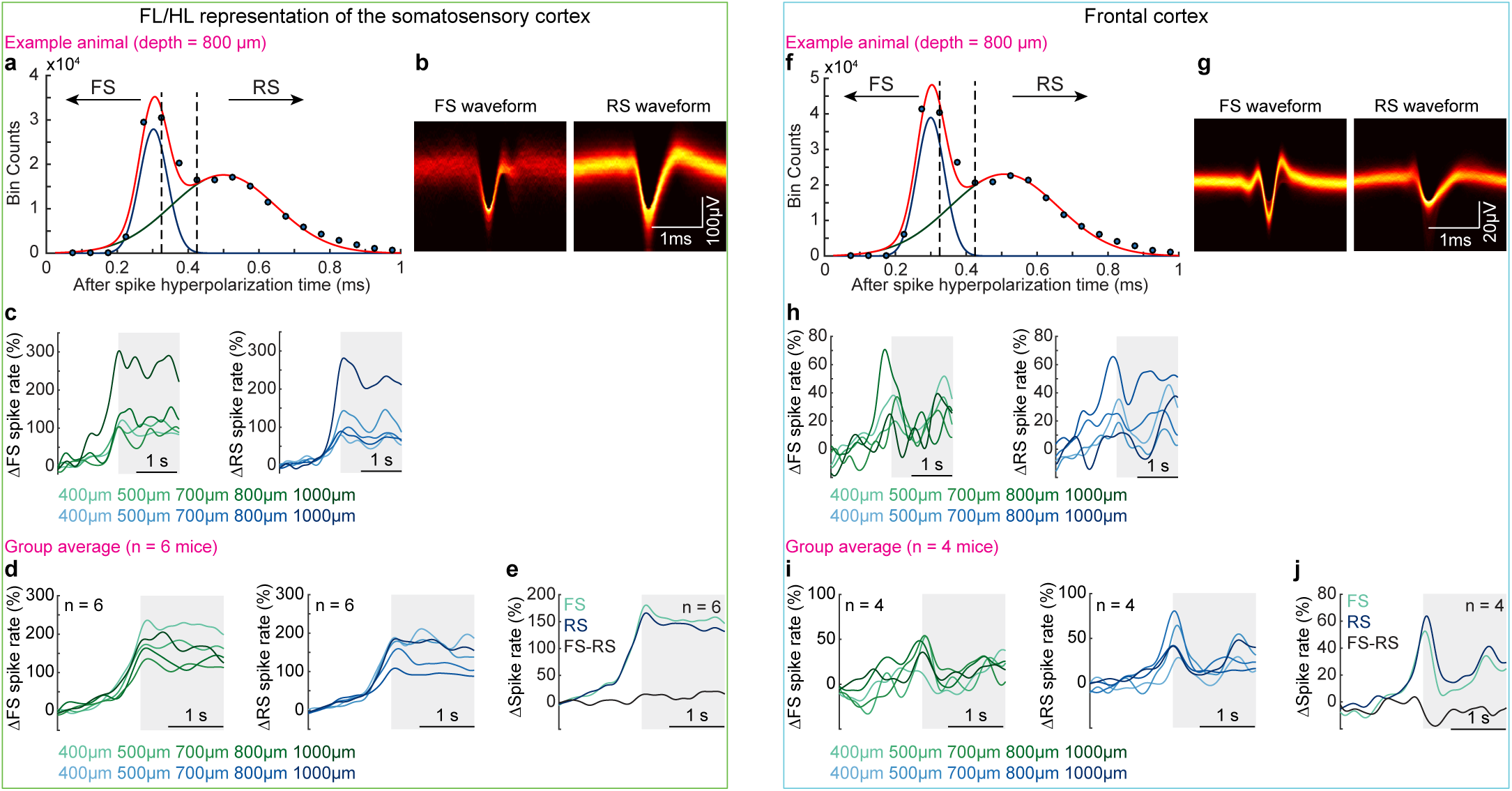
Classification of regular-spiking and fast-spiking neurons based on action potential waveforms and their rate modulations during locomotion in both frontal and somatosensory cortices. Related to **Fig. 1**e-i. We performed spike sorting for neural activity signals acquired using laminar electrodes in both the forelimb/hindlimb representation of the somatosensory cortex (FL/HL, **a-e**) and the frontal cortex (FC, **f-j**). The results shown in (**a**)-(**c**) and (**f**)-(**h**) were from the same mouse shown in **Fig. 1**f, g. (**a**) and (**f**) Histogram of action potential (AP) peak-to-trough durations. (**b**) and (**g**) The waveforms of regular spiking (RS) and fast spiking (FS) neurons for an example animal. (**c**) and (**h**) Locomotion-evoked spike rate changes for fast spiking neurons (ΔFS, left) and regular spiking neurons (ΔRS, right) across different cortical layers for an example animal. Gray shaded area denotes the locomotion period. (**d**) and (**i**) Group average of locomotion-evoked spike rate changes for fast spiking neurons (ΔFS, left) and regular spiking neurons (ΔRS, right) across different cortical layers. Gray shaded area denotes locomotion. (**e**) and (**j**) Group average of locomotion-evoked spike rate changes for FS and RS neurons, as well as the difference between changes of FS and RS spike rates. Gray shaded area denotes locomotion.

**Supplementary Fig. 3.**
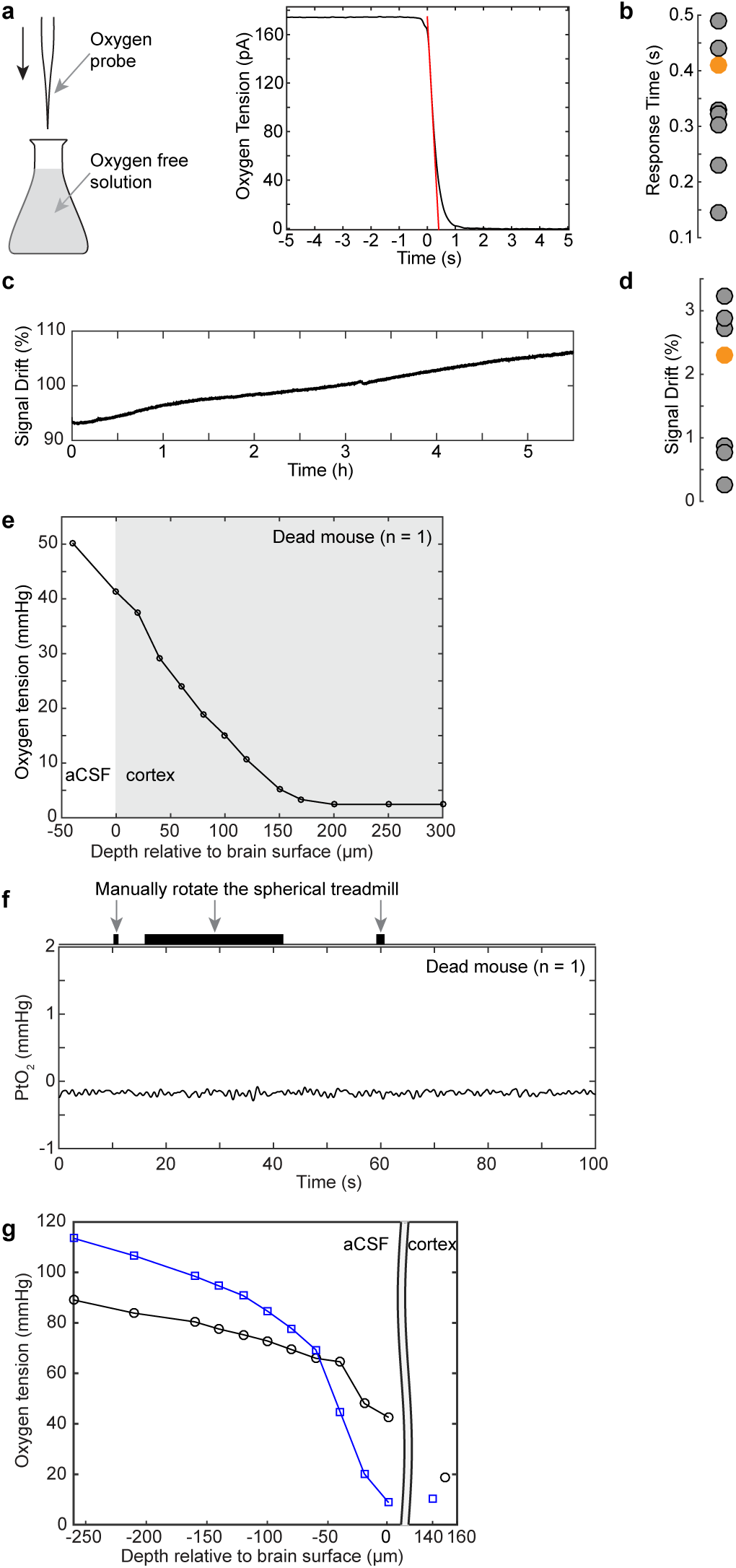
Calibration and properties of oxygen-sensitive electrodes. Related to Fig. 2 and Fig. 3. (**a**) Example trace showing the oxygen electrode signal in response to a step change in oxygen concentration. The oxygen electrode response curve was measured by rapidly immersing the electrode into oxygen-free solution (0.1 M sodium hydrochloride and 0.1 M sodium ascorbate solution). The response time of oxygen electrodes was calculated from this curve. (**b**) Average response time for all electrodes used in this study (n = 9). The orange circle indicates the response time of the probe showing in (**a**). (**c**) Example trace showing the oxygen signal change over a 5-hour time period. The stability of the oxygen electrode was tested by quantifying the signal drift while the electrode was immersed in room temperature water for at least 3 hours. (**d**) Average signal drift for a subset of the electrodes (n = 7, 1.86 ± 1.19% per hour) used in this study. The orange circle indicates the probe showing in (**c**). (**e**) Diffusion of oxygen from the air into the cortex of a dead mouse. To verify the observed oxygen response to locomotion was driven by local perfusion, we also measured oxygen responses in a dead mouse. A similar surgical procedure was applied as described before, and mouse was sacrificed by lung puncture using a 27-gauge needle under deep anesthesia. Oxygen measurements started ∼1 hour after the procedure. The oxygen level in the superficial cortex layers of the dead mouse brain were elevated by the oxygen dissolved in the aCSF bathing the craniotomy. (**f**) An example trace showing the oxygen response in the dead mouse brain to ball rotation at 300 µm below the pia. No noticeable changes of PtO_2_ were observed during manual ball rotation, showing oxygen signal are not due to electrical noise generated by movement. (**g**) Oxygen levels in the aCSF bathing the craniotomy plotted as a function of distance from the pia in two awake mice during rest. Note that oxygen levels drop near the brain due to its metabolic activity.

**Supplementary Fig. 4.**
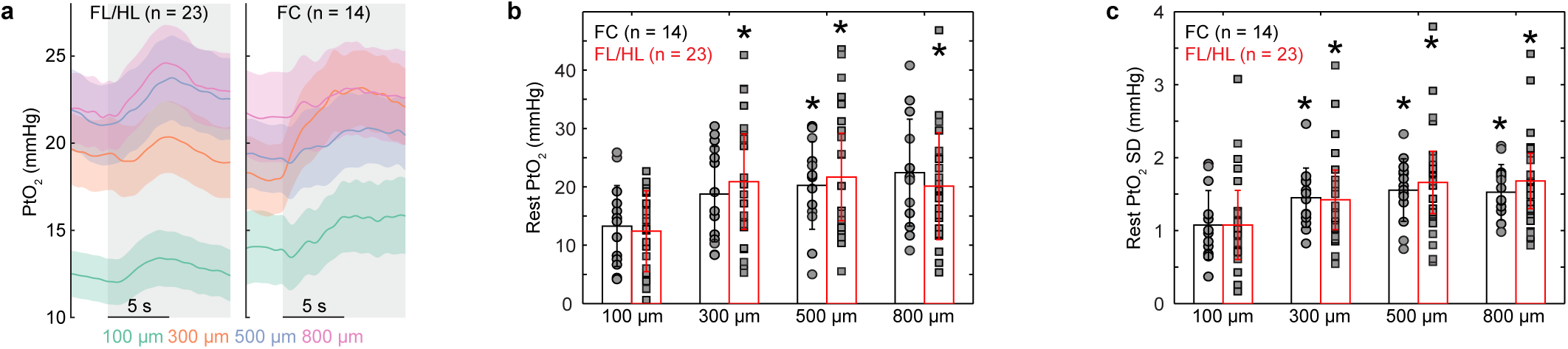
Resting tissue oxygenation is cortical-depth dependent. Related to **Fig. 2**a-d. (a) Locomotion-evoked oxygen increases at all measured depths in both FL/HL (left, n = 23 mice) and FC (right, n = 14 mice). Gray shaded area indicates locomotion. Data are shown as mean ± standard error of the mean (SEM). (**b**) PtO_2_ varies across cortical depth at rest in both the forelimb/hindlimb representation of the somatosensory cortex (FL/HL, n = 23 mice, squares) and the frontal cortex (FC, n = 14 mice, circles). Each square/circle represents data from a single mouse. In FL/HL, resting PtO_2_ was lower at 100 µm (12.41 ± 6.33 mmHg) compared to 300 µm (20.88 ± 10.62 mmHg), 500 µm (21.69 ± 11.29 mmHg) and 800 µm (20.11 ± 9.26 mmHg) below the pia (Kruskal-Wallis ANOVA, F (3, 92) = 11.41, p = 0.0097). In FC, resting PtO_2_ was lower at 100 µm (13.27 ± 6.94 mmHg) compared to 800 µm (22.44 ± 9.17 mmHg, p = 0.0226), but not different from 300 µm (18.77 ± 8.19 mmHg) and 500 µm (20.24 ± 7.50 mmHg) (Kruskal-Wallis ANOVA, F (3, 52) = 8.5, p = 0.0367) below the pia. Resting PtO_2_ were similar at each cortical depth between FL/HL and FC (Mann-Whitney U-test, p > 0.4 for all cortical depths). (c) As in (b) but for the standard deviation (SD) of resting PtO_2_. In FL/HL, SD (Kruskal-Wallis ANOVA, F(3,92) = 14.7, p = 0.0021) was smaller at 100 µm (1.08 ± 0.68 mmHg) compared to 500 µm (1.66 ± 0.75 mmHg, p = 0.0103) and 800 µm (1.68 ± 0.65 mmHg, p = 0.0034), but not 300 µm (1.42 ± 0.66 mmHg) below the pia. In FC, the SD (Kruskal-Wallis ANOVA, F(3, 52) = 8.99, p = 0.0294) was smaller at 100 µm (1.08 ± 0.47 mmHg) compared to 500 µm (1.55 ± 0.43 mmHg, p = 0.0254) and 800 µm (1.53 ± 0.38 mmHg, p = 0.0418), but not layer 300 µm (1.45 ± 0.41 mmHg) below the pia. Resting fluctuations of PtO_2_ were similar at each cortical depth between FL/HL and FC (Mann-Whitney U-test, p > 0.4 for all cortical depths).

**Supplementary Fig. 5.**
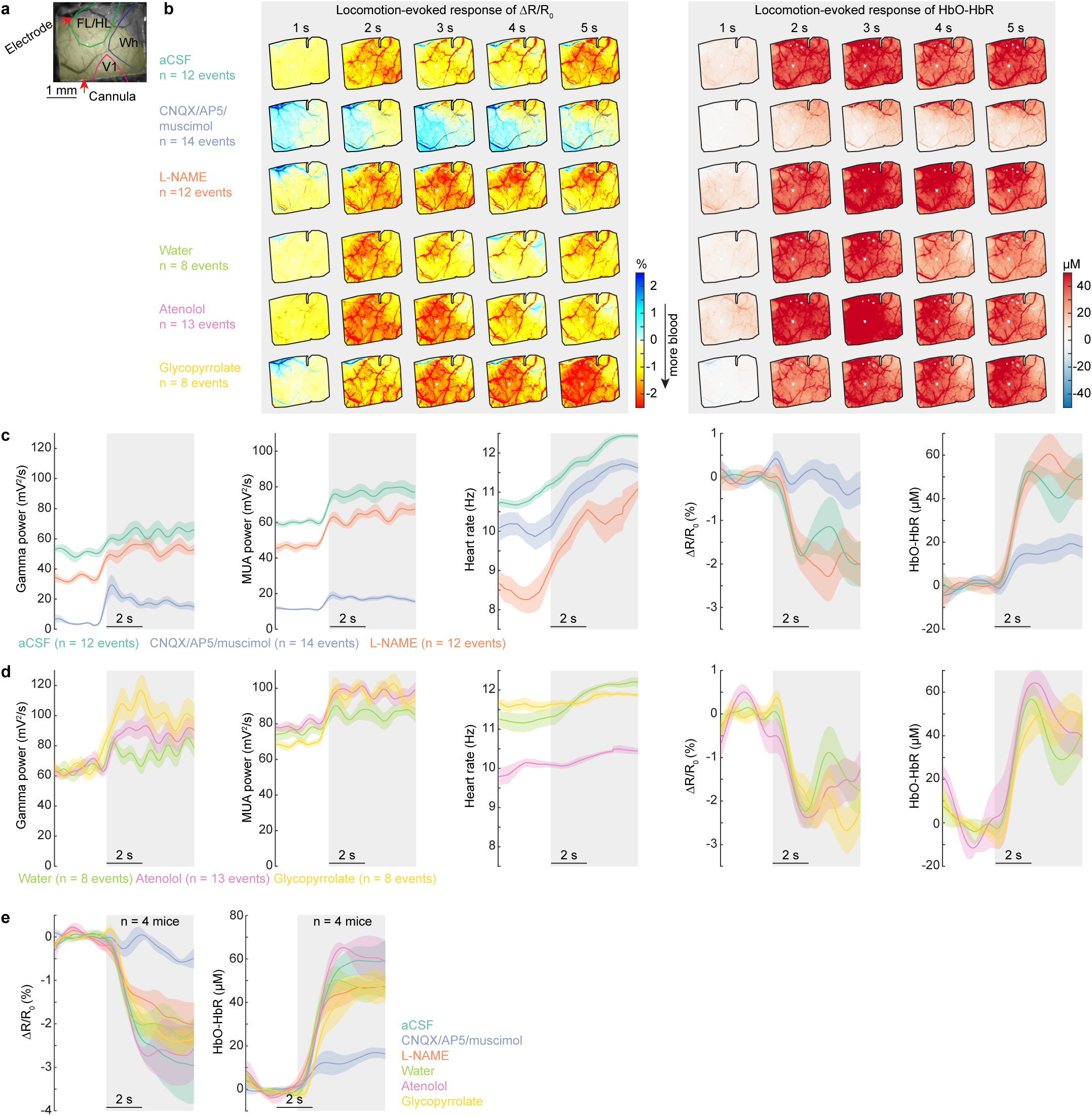
Locomotion-evoked hemodynamic responses depend on local neural activity, not cardiovascular responses. Related to Fig. 3a-d. (**a**) An image of a polished thin-skull window with cannula and electrode implants. The yellow shaded area indicates the area affected by the drug infusion as determined from electrical recording. This area is used as the ROI for quantification of hemodynamic signals. (**b**) Locomotion-evoked spatial distribution of ΔR/R_0_ (left) and HbO-HbR (right) for the same mouse shown in Fig. 3a-c, following intracerebral infusion of aCSF (n = 12 locomotion events), CNQX/AP5/muscimol (n = 14 locomotion events) and L-NAME (n = 12 locomotion events), as well as intraparietal injection of water (n = 8 locomotion events), atenolol (n = 13 locomotion events) and glycopyrrolate (n = 8 locomotion events). (**c**) Locomotion-evoked response of gamma-band power, MUA power, heart rate, ΔR/R_0_ and HbO-HbR following intracerebral infusion of aCSF, CNQX/AP5/muscimol and L-NAME in the same animal shown in (**b**). (**d**) As (**c**) but for responses following intraparietal injection of water, atenolol and glycopyrrolate. (**e**) Group average (n = 4 mice) of locomotion-evoked ΔR/R_0_ (left) and HbO-HbR (right) following different drug administration. Note that only disruption of neural activity (CNQX/AP5/muscimol infusion) caused changes in locomotion-evoked vasodilation or oxygenation, while glycopyrrolate and atenolol had large effects on heart rate. This shows that the oxygenation responses observed here are not affected by cardiovascular changes.

**Supplementary Fig. 6.**
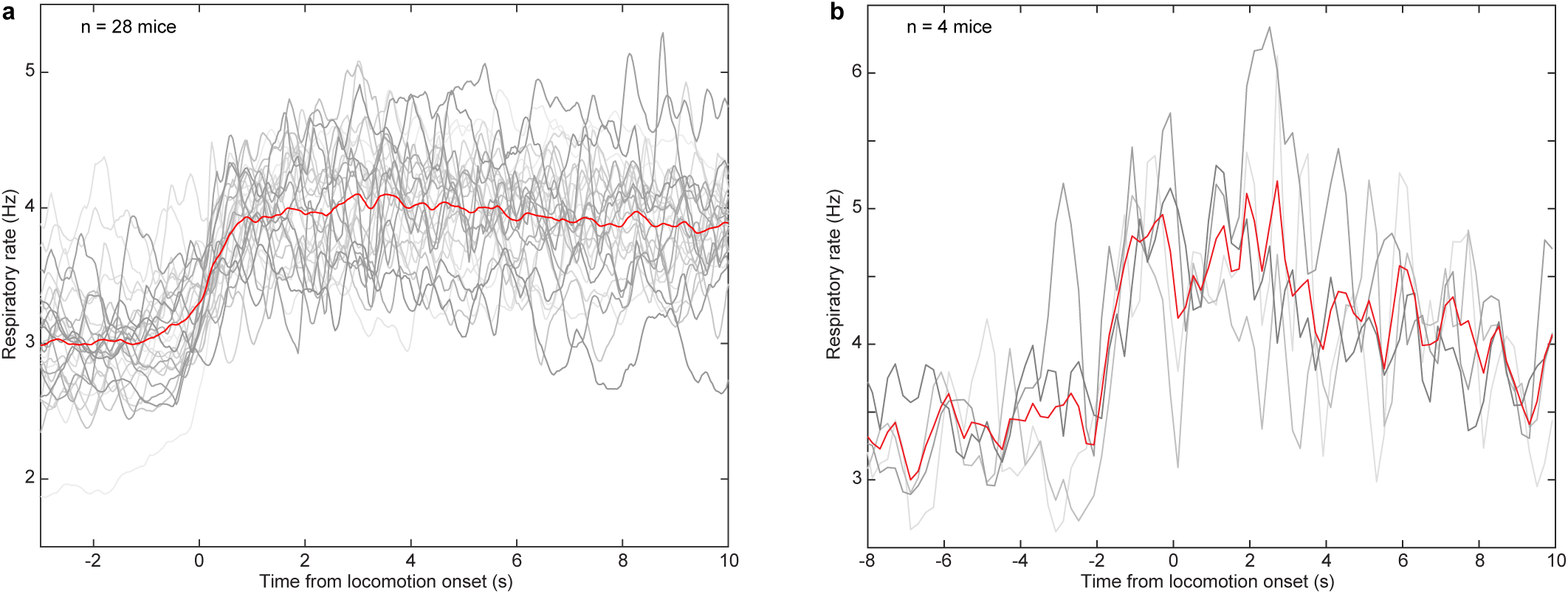
Measuring respiration with a thermocouple. Related to Fig. 4. (**a**) Respiratory rate at rest and during locomotion (n = 28 mice) for mice running on the spherical treadmill. Each gray trace indicates the averaged respiratory rate from one mouse (∼ 2 h recording), and the red trace indicates group average. Time zero indicates onset of locomotion. (**b**) As in (**a**) but for mice (n = 4) running on the rotating disk.

**Supplementary Fig. 7.**
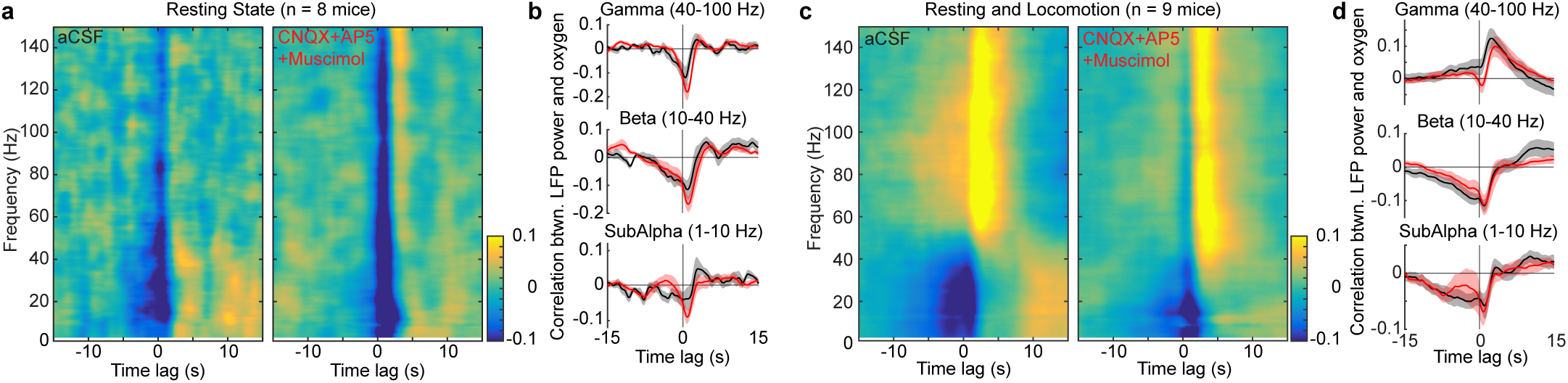
Suppressing vasodilation does not change the correlation between oxygenation and neural activity. Related to Fig. 4a-e. (**a**) Group average (n = 8 mice) of cross-correlation between PtO_2_ and LFP at various frequency band during periods of rest after aCSF (left) and CNQX/AP5/muscimol (right) administration. (**b**) Cross-correlation between PtO_2_ and LFP at different frequency band during periods of rest after aCSF (black) and CNQX/AP5/muscimol (red) administration. The shaded region shows the population standard error of the mean (n = 8 mice). (**c**) As in (**a**) but for periods including both rest and locomotion (n = 9 mice). (**d**) As in (**b**) but for periods of both rest and locomotion (n = 9 mice).

**Supplementary Fig. 8.**
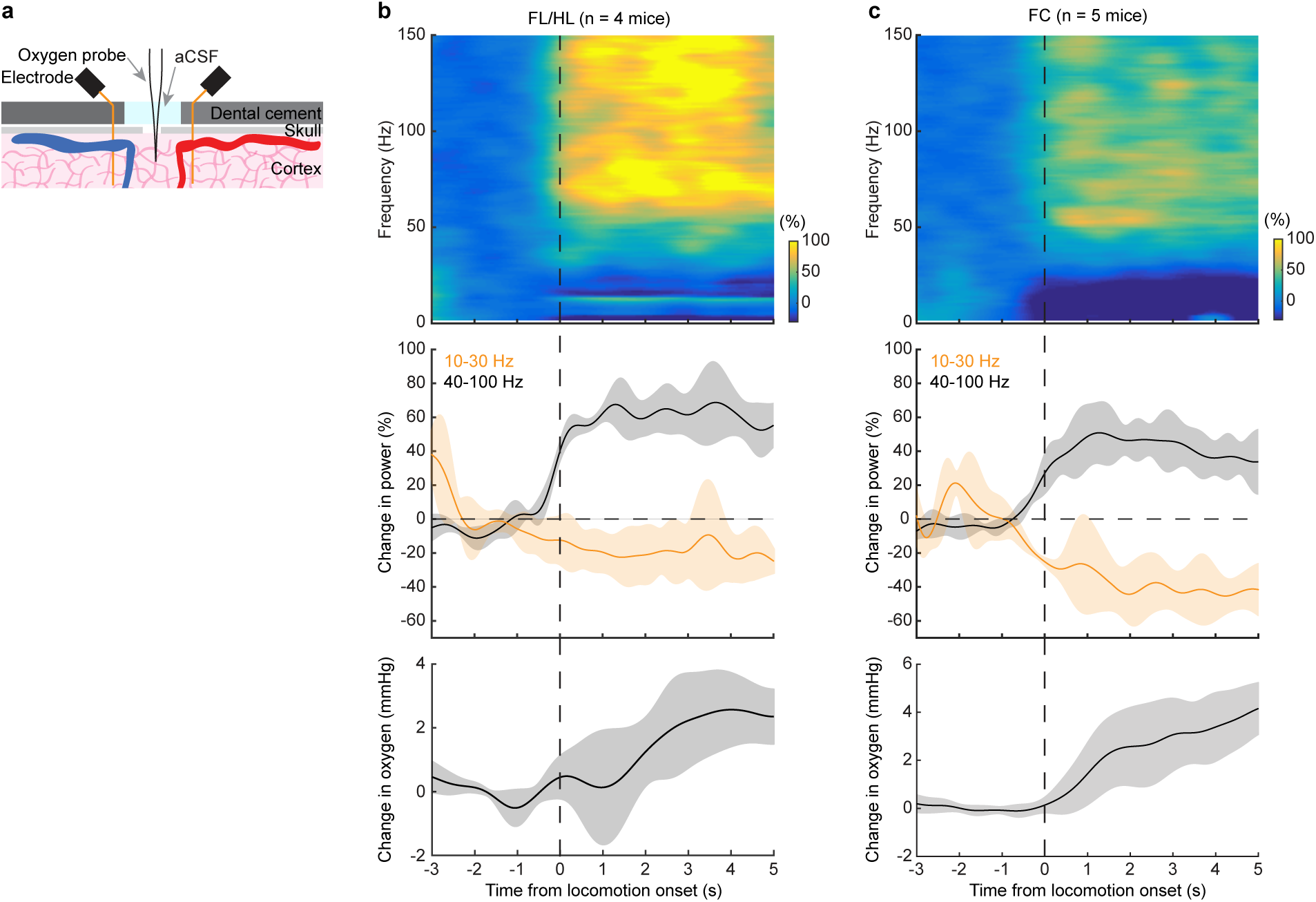
Locomotion-evoked LFP power changes in both FL/HL and FC. (**a**) Schematic of experimental setup for simultaneous tissue oxygenation and electrophysiology measurement. (**b**) Top, time-frequency representation of locomotion-evoked changes in LFP power in FL/HL (n = 4 mice). Middle, locomotion-evoked changes of gamma-band (40-100 Hz, black) and beta-band (10-30 Hz, orange) power. Data were shown as mean ± SD. Bottom, locomotion-evoked changes of tissue oxygenation. Data were shown as mean ± SD. (**c**) As (**b**) but for FC (n = 5 mice). The data used in (**b**) and (**c**) were from the same group of mice shown in Fig. 3g-h with the craniotomy superfused with aCSF.

**Supplementary Fig. 9.**
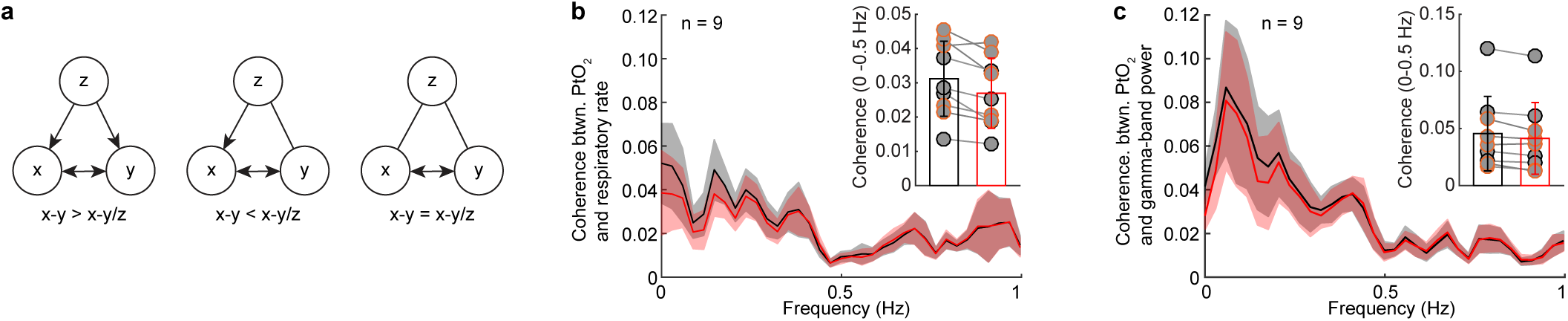
Respiration and neural activity modulate tissue oxygenation independently. Related to Fig. 4. (**a**) Schematic showing different patterns of coupling between three signals (x, y and z) that can be revealed by the partialization technique. Partialization with z (x-y/z) may decrease the x-y coherence if they are both connected to z, or even completely block the coherence if they are exclusively drive by signal z (left). Partialization with z may increase the x-y coherence if x and y are affected by z in an asymmetric manner (middle). Partialization with z may not affect the x-y coherence if x and y are not connected to z (right). (**b**) Group average of coherence between respiratory rate and PtO_2_ before (black) and after (red) partializing the effect of neural activity. Shaded area denotes mean ± SE. The inset denotes that group average of coherence within the frequency range of 0 to 0.5 Hz before (black) and after (red) partialization. (**c**) As in (**b**) but for coherence between gamma-band LFP power and PtO_2_before (black) and after (red) partializing the effect of respiratory rate.

